# Ankrd26 is critical for cell differentiation and cancer-linked mutations affect its key properties

**DOI:** 10.1101/2021.05.19.444897

**Authors:** Sarah A. Hofbrucker-MacKenzie, A. Sofie Englisch, Maryam Izadi, Klara Metzner, Michael M. Kessels, Britta Qualmann

## Abstract

Derailed signaling originating from the plasma membrane is associated with many types of cancer. Different human cancers and thrombocytopenia are linked to *ANKRD26* mutations. We unveil that Ankrd26 is a plasma membrane-localized protein forming nanoclusters and that Ankrd26 is critical for retinoic acid/BDNF-induced neuroblastoma differentiation. An N-terminal amphipathic structure lacking in an AML-associated Ankrd26 mutant is indispensable for membrane binding and bending by partial membrane insertion and renders Ankrd26 inactive in both gain-of-function and loss-of-function/rescue studies addressing cellular differentiation. In a papillary thyroid carcinoma-linked mutant, truncated Ankrd26 is fused with the kinase domain of the protooncogene RET. Our data show that the Ankrd26 part of this fusion mutant mediates anchoring of the RET kinase domain to the plasma membrane and self-association by the coiled coil domain of Ankrd26. Ankrd26-RET fusion led to massively increased ERK1/2 activity and RET autophosphorylation at both Y905 and Y1015, i.e. caused aberrant RET signaling. Our results highlight the importance and molecular details of Ankrd26-mediated organizational platforms for cellular differentiation and signaling pathways from the plasma membrane, which, if derailed, lead to cancer-associated pathomechanisms involving the unveiled Ankrd26 properties.

## Introduction

The plasma membrane represents the cellular interface to the outer world and thus transmits and integrates a plethora of signals towards the cytoplasm. Plasma membrane-originating signal transduction pathways in turn also modulate the organization and shape of the plasma membrane when cells adapt and respond to such signals. Derailed signaling pathways originating from the plasma membrane are associated with many types of cancer. Thus, proteins linked to signaling pathways that might furthermore spatially organize signaling-competent membrane domains and/or modulate the topology of membranes take center stage in contemporary research.

Several, yet molecularly seemingly very distinct mutations in *ANKRD26* have been linked to different human cancers (Cerami et al., 2012; Marconi et al., 2017; Staubitz et al., 2019) and also to thrombocytopenia (Noris et al., 2011; Pippucci et al., 2011; Bluteau et al., 2014; Wahlster et al., 2021). These mutations include a deletion of a more C terminal part of Ankrd26 leading to a disruption of Ankrd26’s binding to PIDD1 – a component of a multi-protein complex (PIDDosome) involved in the cellular response to extra centrosomes (Cerami et al., 2012; Fava et al., 2017; Evans et al., 2021; Burigotto et al., 2021). More recently, an N-terminal truncation has been found to be associated with acute myeloid leukemia (AML) (Marconi et al., 2017). Furthermore, the cancer-linked Ankrd26 mutations include a C terminal Ankrd26 truncation and fusion with a part of the protein product of the protooncogene *RET* (*rearranged during transfection*) (Staubitz et al., 2019) – the tyrosine kinase signaling component of multisubunit receptor complexes for glia cell line-derived neurotrophic factor (GDNF) family ligands (GFLs), which frequently shows partial recombination with other genes in a variety of cancers (Shaw et al., 2013; Liu et al., 2021).

The cancer-associated Ankrd26 mutations found up to today are molecularly heterogeneous and therefore do not provide an easy avenue to understanding Ankrd26-associated diseases. Similarly, although Ankrd26 was first described as protein with similarities to POTEs (expressed in prostate, ovary, testis, and placenta) (Bera et al., 2002; Bera et al., 2008), which are considered as potential biomarkers and therapeutic targets based on their association with poor prognosis in ovarian and other cancers (Bera et al., 2006; Redfield et al., 2013), the comparison of Ankrd26 to POTE proteins is unrewarding. First, also functions and molecular properties of POTEs are largely unknown. Second, the suggested similarity of POTEs to Ankrd26 is in fact very limited. Although both contain ankyrin repeats, they reside in the middle in POTEs (Bera et al., 2002) but in the N terminal part in Ankrd26. Also, Ankrd26 lacks the cysteine-rich domain typical for POTEs (Bera et al., 2002). Due to this lack of knowledge about the properties and molecular characteristics of Ankrd26, Ankrd26 dysfunctions in pathophysiological processes in human patients largely remained elusive.

We unveil that Ankrd26 is a partially clustered, plasma membrane-localized protein that has the ability to shape membranes by employing an N terminal, membrane-inserted amphipathic structure and a membrane curvature-sensing ankyrin repeat array. Furthermore, Ankrd26 is able to form larger, membrane-associated arrays by self-association. Ankrd26 loss-of-function studies unveiled a strong impairment in the morphological differentiation of neuroblastoma cells. Its properties and their underlying molecular mechanisms demonstrate that Ankrd26 is a member of the recently suggested N-Ank protein superfamily (Wolf et al., 2019). The AML-associated mutation of Ankrd26 disrupted the membrane binding and shaping capability of Ankrd26 and its functions in cellular differentiation. The cancer-associated RET fusion led to a protein that is fully self-association-competent, is constitutively targeted to the plasma membrane and results in aberrant RET signaling by a massive increase in two forms of RET autophosphorylation as well as to activation of ERK1/2. Understanding the cellular functions and molecular mechanistic properties of Ankrd26 significantly advances our understanding of Ankrd26-associated pathomechanisms in human cancer patients.

## Results

### Ankrd26 associates with the plasma membrane using an N-Ank module

Ankrd26 is involved in several types of cancer (Cerami et al., 2012; Marconi et al., 2017; Staubitz et al., 2019). Yet, the molecular properties and the functions of Ankrd26 largely remained elusive. Ankrd26-associated thrombocytopenia is caused by 5’UTR mutations that seem to lead to some N terminal truncation. Also an AML-associated mutation of Ankrd26 is marked by an N-terminal truncation of Ankrd26 (Marconi et al., 2017). These findings urgently call for thus far lacking in-depth-analyses of Ankrd26’s N terminal part to understand the underlying pathomechanisms.

Although Ankrd26 has been described as cytosolic protein in neurons and glia cells (Acs et al., 2015) and more recent studies described it as centriolar protein despite the fact that neither ciliogenesis nor centriole duplication was disrupted in cells lacking Ankrd26 (Yan et al., 2020; Evans et al., 2021; Burigotto et al., 2021), our subcellular fractionation studies unveiled a strong membrane association. Similar to farnesylated mCherry (CherryF) coexpressed as marker for constitutively plasma membrane-associated proteins, a significant Ankrd26 portion was detected in the fractions P2 and P2’ (**Figure 1A,B**).

**Figure 1.**
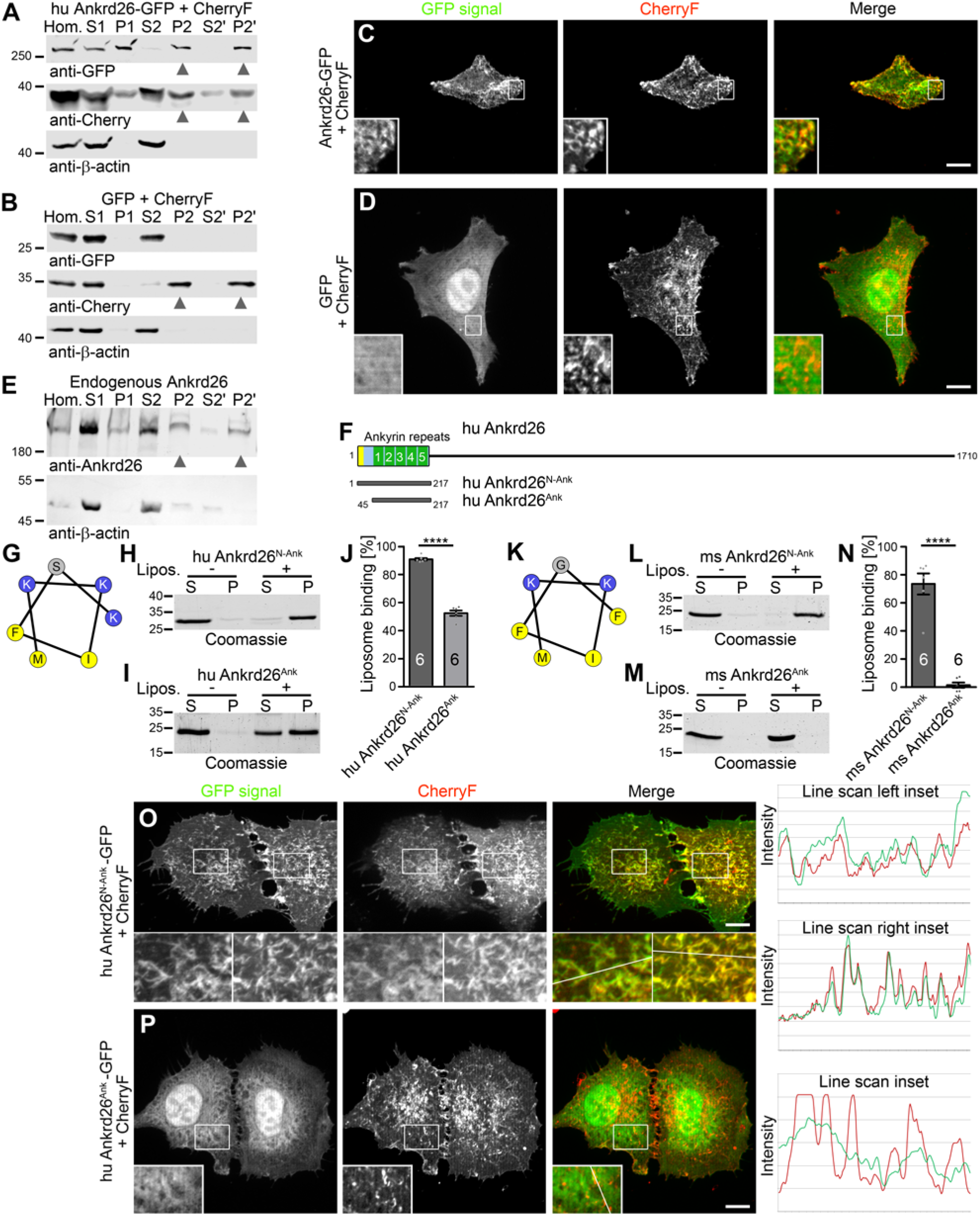
Ankrd26 is a plasma membrane-binding protein using a putative N-Ank module for membrane association. **(A,B)** Subcellular fractionations of human (hu) Ankrd26-GFP- and GFP-transfected HEK293 cells coexpressing CherryF. Arrowheads highlight proteins detected in P2 and P2’ membrane fractions. CherryF represents plasma membrane-integrated proteins; endogenous β-actin represents cell cortex-associated and cytosolic proteins. **(C,D)** Maximum intensity projections (MIPs) of Ankrd26-GFP and GFP in HeLa cells cotransfected with CherryF. Insets represent magnifications of boxed areas. See **Figure 1-figure supplement 1** for additional intensity plots of the red and green fluorescent channel along the randomly positioned lines. Bars, 10 µm. **(E)** Anti-Ankrd26 and anti-β-actin immunoblottings of a subcellular fractionation of HEK293 cells. Arrowheads highlight endogenous Ankrd26 in membrane fractions P2 and P2’. **(F)** Schematic representation of human Ankrd26 (UniProt Q9UPS8) with its apparent N terminal N-Ank module containing five ankyrin repeats (UniProt; see **Figure 1-figure supplement 1** for comparison of all suggested repeats with a general ankyrin repeat consensus). **(G-N)** *In silico* analyses of putative amphipathic structures at the N termini of human **(G)** and mouse (ms) Ankrd26 **(K)** (helical wheel representations using HELIQUEST; amino acid color coding: blue, positively charged; yellow, hydrophobic; grey, other) and representative example images of Coomassie-stained SDS-PAGE analyses of liposome coprecipitation studies (S, supernatant; P, pellet) addressing a membrane binding of recombinant purified human and mouse Ankrd26 (UniProt D3Z482) N-Ank modules (Ankrd26^N-Ank^) and deletion mutants thereof comprising only the ankyrin repeat arrays (Ankrd26^Ank^), i.e. lacking the putative amphipathic structures at the N termini **(H,I,L,M)** as well as corresponding quantitative analyses of liposome binding **(J,N)**. Data represent mean±standard error of means (SEM) of quantitative analyses with two independent liposome preparations (n=6 experiments each condition). Shown are bar plots with individual data points (dot plots). Two-tailed, unpaired Student’s t-test. \*\*\*\**P*<0.0001. **(O,P)** Analysis of the localization of human Ankrd26^N-Ank^ **(O)** and of the Ankrd26^Ank^ deletion mutant **(P)** in HeLa cells coexpressing CherryF. Shown are MIPs and line scans along lines depicted in the inset representing magnifications of boxed areas. For GFP control and related analyses with murine proteins see **Figure 1-figure supplement 2.** Bars, 10 µm.

Also in intact cells, Ankrd26-GFP specifically localized to CherryF-marked folds of the plasma membrane (**Figure 1C,D; Figure 1-figure supplement 1A,B**). Importantly, membrane association was also demonstrated for endogenous Ankrd26, which clearly was detectable in fractions P2 and P2’ (**Figure 1E**).

Sequence analyses and structure predictions of human Ankrd26 unveiled five N terminal ankyrin repeats between amino acid 46 and 207 and an N terminal amphipathic structure, which together may represent an N-Ank module (**Figure 1F,G**; **Figure 1-figure supplement 1C,D**). Indeed, purified, recombinant human Ankrd26^1-217^ (hu Ankrd26^N-Ank^) coprecipitated with liposomes in *in vitro* reconstitutions (**Figure 1H**). In contrast, an N terminal deletion mutant of Ankrd26 lacking the suggested amphipathic structure (hu Ankrd26^Ank^, hu Ankrd^45-217^) was significantly impaired in liposome binding (**Figure 1I,J**). The N terminus of Ankrd26 thus is important for Ankrd26’s membrane association. Similar results were obtained for the mouse Ankrd26 N-Ank module (ms Ankrd26^N-Ank^, ms Ankrd26^1-208^) and the respective deletion mutant (ms Ankrd26^Ank^, ms Ankrd26^11-208^). Deletion of the N terminus suppressed the membrane binding almost completely (**Figure 1K-N**). Also in cells, both mouse and human Ankrd26 N-Ank modules clearly colocalized with CherryF at the plasma membrane, whereas both corresponding N terminal deletion mutants failed to show any specific membrane localization (**Figure 1O,P**; **Figure 1-figure supplement 2)**.

Ankrd26 thus is a membrane-binding protein, associates with membranes directly, as proven by *in vitro* reconstitution using purified components, and this ability is brought about by an N terminal domain that may resemble an N-Ank module.

### A mutated Ankrd26 N-Ank domain found in AML patients is deficient for membrane binding

Our data suggested that an *ANKRD26* mutation found in AML patients, which gives rise to an N terminal deletion (Marconi et al., 2017), is likely to represent a loss-of-function mutation for Ankrd26’s N-Ank-mediated membrane binding (**Figure 2A**). Indeed, the disease mutation identified in AML rendered the N-Ank module (hu Ankrd26^N-Ank^°^AML^; Ankrd26^78-217^) completely unable to associate with membranes in *in vitro* reconstitutions (**Figure 2B-D**). In fractionation studies, Ankrd26^N-Ank^°^AML^ was absent from the membrane fractions P2 and P2’, whereas the corresponding WT protein (Ankrd26^N-Ank^) was detectable in both P2 and P2’ (**Figure 2E,F**). Consistently, immunofluorescence analyses demonstrated that Ankrd26^N-Ank^°^AML^ failed to properly localize to the plasma membrane. Instead, Ankrd26^N-Ank^°^AML^ was distributed aberrantly in the cytosol (**Figure 2G**). Importantly and in line with a critical role of the N terminal helix of the proposed N-Ank module for membrane association, AML mutation in the full-length protein context (hu Ankrd26°^AML^; Ankrd26^78-1710^) led to a protein that was unable to bind to membranes. Both membrane fractions, P2 and P2’, were devoid of the AML mutant of full-length Ankrd26 (**Figure 2H**). Thus, Ankrd26’s involvement in AML seems to be related to a defective membrane targeting of the Ankrd26 mutant found in AML patients.

**Figure 2.**
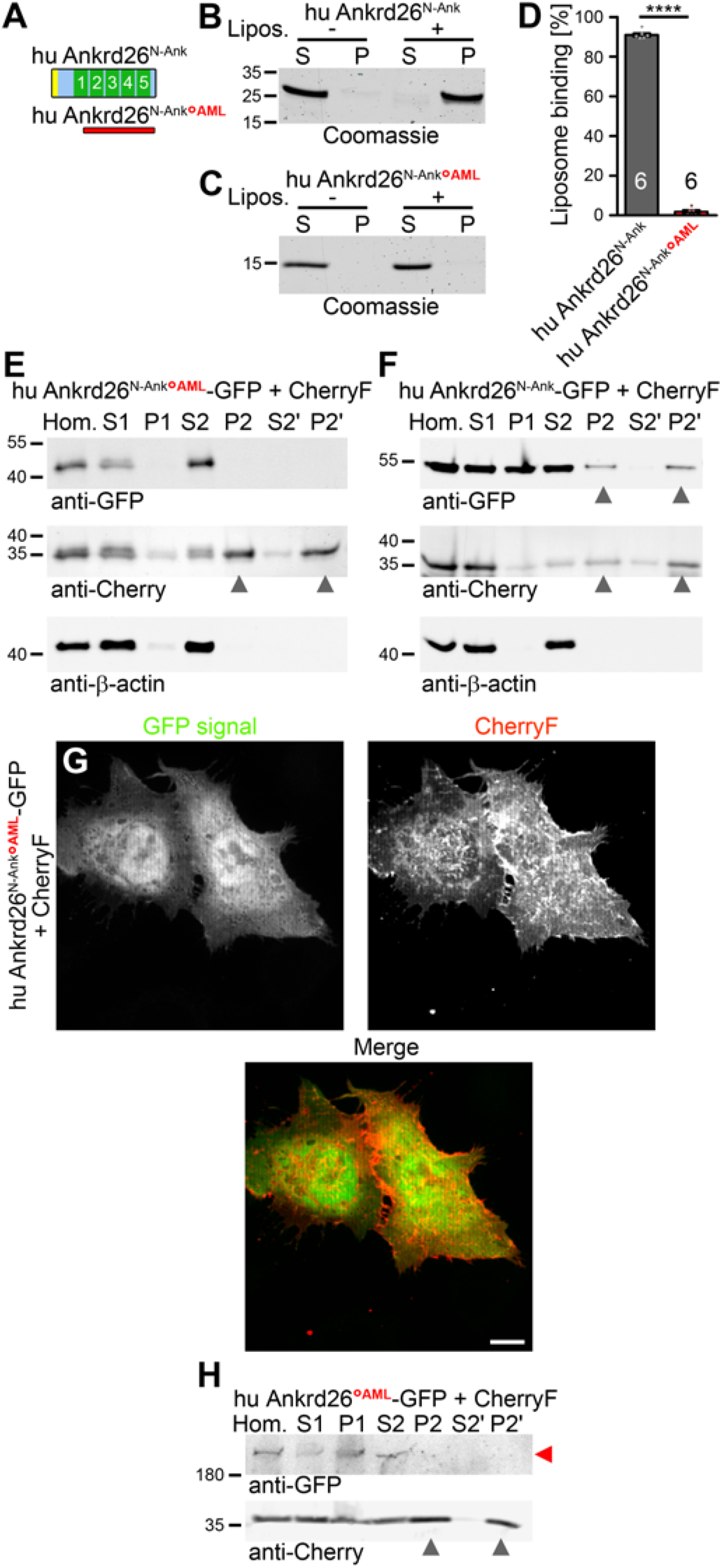
An N terminal truncation of the suggested N-Ank module of Ankrd26 found in AML patients completely disrupts membrane binding *in vitro* and *in vivo*. **(A)** Schematic representation of the ankyrin repeats-containing N termini of human Ankrd26 and of a corresponding mutant found in AML patients, which results in an N terminally truncated protein (Ankrd26^78-217^; Ankrd26^N-Ank^°^AML^). **(B-D)** Representative images of Coomassie-stained gels of liposome coprecipitation analyses with Ankrd26^N-Ank^ and Ankrd26^N-Ank^°^AML^ (side-by-side analyses using the same preparation of liposomes) **(B,C)** and quantitative liposome binding evaluations thereof **(D)**. **(E,F)** Immunoblotting analyses of subcellular fractionations of lysates of HEK293 cells that were transfected with Ankrd26^N-Ank^°^AML^-GFP **(E)** and Ankrd26^N-Ank^-GFP **(F)**, respectively, and coexpressed CherryF. Arrowheads highlight proteins detected in P2 and P2’ membrane fractions (WT Ankrd26^N-Ank^ and CherryF only). **(G)** MIP showing that Ankrd26^N-Ank^°^AML^-GFP did not localize to the plasma membrane of HeLa cells marked by coexpressed CherryF. **(H)** Immunoblotting analyses of subcellular fractionations of lysates of HEK293 cells that were transfected with Ankrd26°^AML^-GFP and CherryF. Grey arrowheads highlight P2 and P2’ membrane fractions (containing the plasma membrane marker CherryF but being devoid of Ankrd26°^AML^-GFP (red arrowhead)). Bar, 10 µm. Data, mean±SEM. Quantitative analyses with two independent liposome preparations (n=6 experiments each condition) **(D)**. Bar plots with individual data points (dot plots). Two-tailed, unpaired Student’s t-test. \*\*\*\**P*<0.0001.

### The part lacking in the Ankrd26 AML mutant responsible for Ankrd26’s tight membrane binding comprises an amphipathic helix

We next addressed the mechanisms of Ankrd26’s membrane association disrupted by the N terminal deletion in the Ankrd26 AML mutant. An N terminal amphipathic structure may mediate integration into the membrane instead of mere electrostatic surface association. Indeed, the WT Ankrd26 N-Ank module was not sensitive to suppressing electrostatic interactions by rising salt concentrations but showed full salt resistance even at 250 mM NaCl (**Figure 3A,B**). Similarly, also the membrane-binding of mouse Ankrd26^N-Ank^ was resistant to suppression of electrostatic interactions (**Figure 3- figure supplement 1**). In contrast, the (reduced) membrane binding of Ankrd26^Ank^ (compare also **Figure 1I,J**) was sensitive to rising salt concentrations and therefore apparently predominantly based on electrostatic interactions (**Figure 3C,D**).

**Figure 3.**
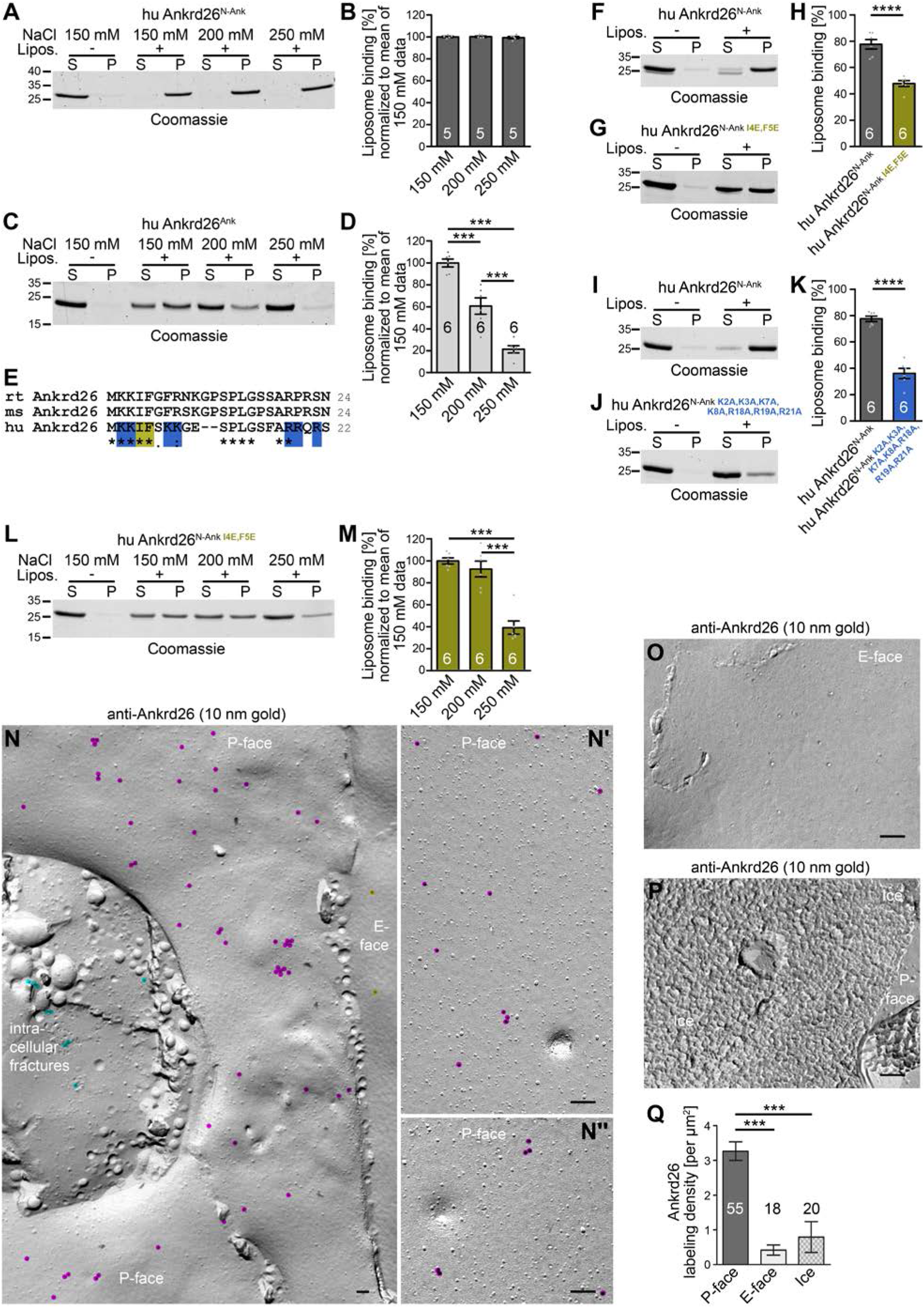
Ankrd26 is a plasma membrane-localized and –embedded N-Ank protein. **(A-D)** Representative images of Coomassie-stained SDS-PAGE gels **(A,C)** of attempts of extraction of liposome-bound human Ankrd26^N-Ank^ **(A,B)** and the corresponding Ankrd26^Ank^ deletion mutant **(C,D)** by increasing salt concentrations (S, supernatant; P, pellet) and quantitative analyses thereof showing that the liposome binding of Ankrd26^N-Ank^ is fully salt-resistant, whereas the weaker membrane association of Ankrd26^Ank^ was merely based on electrostatic interactions suppressible by increasing salt **(B,D)**. **(E)** Alignment of the N termini of rat (rt; UniProt, M0R3T8), mouse (ms; UniProt, D3Z482) and human (hu; UniProt, Q9UPS8) Ankrd26 with a high conservation of both hydrophobic and positively charged amino acids. Positively charged residues are highlighted in blue and mutated hydrophobic residues are highlighted in dark yellow. **(F-K)** Liposome binding analyses with mutant N-Ank modules of Ankrd26 with either two hydrophobic residues turned into hydrophilic ones (Ankrd26^N-Ank^ ^I4E,F5E^) and with all positive residues of the Ankrd26 N terminal sequence erased (Ankrd26^N-Ank^ ^K2A,K3A,K7A,K8A,R18A,R19A,R21A^), respectively. **(L,M)** Salt extraction trials with Ankrd26^N-Ank^ ^I4E,F5E^ demonstrating a disruption of the salt resistance of the membrane binding of Ankrd26’s N- Ank module by mutating isoleucine 4 and phenylalanine 5. **(N-N’’)** Transmission electron microscopical images of freeze-fractured SK-N-SH cells that were differentiated with retinoic acid/BDNF and immunolabeled with anti-Ankrd26 antibodies (immunolabeling, 10 nm gold) at low **(N)** and high magnification **(N’,N’’)**. Bars, 100 nm. Gold particles were overlayed with enlarged colored dots to improve visibility (magenta, anti-Ankrd26 labeling at the P-face of plasma membrane (cytosolic face); turquoise, anti-Ankrd26 labeling at intracellular membranes; orange, (background) labeling at E-face). **(O,P)** Representative images of E-face membrane areas and ice surfaces devoid of anti-Ankrd26 immunogold labeling. Bars, 100 nm. **(Q)** Quantitative analysis of Ankrd26 labeling densities at P-face areas, E-faces areas and ice. Data represent mean±SEM. **(B,D,H,K,M)** Bar plots with individual data points (dot plots). **(B)** n=5 assays; **(D,H,K,M)** n=6 assays each. **(Q)** n=55 (P-face), 18 (E-face) and 20 (ice) of 1085 µm^2^ (P-face), 240 µm^2^ (E-face) and 62 µm^2^ (ice) total area. **(B,D,M,Q)** one-way ANOVA with Tukey post hoc test; **(H,K)** two-tailed, unpaired Student’s t-test. \*\*\**P*<0.001; \*\*\*\**P*<0.0001.

The amphipathic nature of the N terminal key element for Ankrd26’s tight membrane association was then proven by two sets of mutations erasing on one side two fully conserved hydrophobic residues (I4E, F5E) and on the other side seven hydrophilic, positively charged residues (**Figure 3E**). Both mutants showed significantly reduced membrane binding in quantitative studies (**Figure 3F-K**). Importantly, mutation of the two hydrophobic residues isoleucine 4 and phenylalanine 5 to hydrophilic amino acids (glutamate) was sufficient to render Ankrd26 susceptible to electrostatic suppression of membrane association (**Figure 3L,M**). It can therefore be concluded that the ability of Ankrd26^N-Ank^ to tightly bind to membranes relies on hydrophobic membrane associations and embedding into the hydrophobic phase of the membrane mediated by an amphipathic structure at the N terminus of Ankrd26.

### Endogenous Ankrd26 resides inside the cytosolic leaflet of freeze-fractured plasma membranes

Partial membrane insertion as molecular mechanism of Ankrd26’s membrane binding suggested that it might be possible to establish a detection of endogenous Ankrd26 in cellular membranes by freeze-fracturing, immunogold labeling and transmission electron microscopy (TEM) similar to methods, which we were able to successfully establish for other membrane-inserted proteins, such as caveolins (Koch et al., 2012; Seemann et al., 2017), syndapins (Schneider et al., 2014; Seemann et al., 2017; Izadi et al., 2021), and the N-Ank protein ankycorbin (Wolf et al., 2019). Freeze-fractured plasma membranes of SK-N-SH neuroblastoma cells differentiated with retinoic acid and BDNF according to Encinas et al. (2000) were successfully immunogold labeled with antibodies against Ankrd26 (**Figure 3N**). The labeling was specific, as control surfaces in the same samples, i.e. the E-face of the membrane and ice surfaces, were almost devoid of immunolabeling (**Figure 3O,P**). Quantitative analyses of labeling densities over wide ranges of systematically imaged areas confirmed the specificity of the anti-Ankrd26 immunolabeling at the plasma membrane of neuroblastoma cells (**Figure 3Q**).

Ankrd26 was present across wide areas of the plasma membrane and did not show correlation with any membrane protein complexes already structurally visible by platinum shadowing (**Figure 3N**). Also, anti-Ankrd26 immunolabeling was not correlated with the rare sites of circular membrane invagination of different depth we observed (**Figure 3N’,N’’**) but occurred at various areas of the membrane (**Figure 3N**).

Besides at the plasma membrane, also a rare case of intracellular fracture showed some, albeit low anti-Ankrd26 labeling suggesting that Ankrd26 functions may not exclusively be reflected by the plasma membrane-anchored Ankrd26 prominently observed in our electron microscopical examinations (**Figure 3N-N’’**).

### Ankrd26’s ankyrin repeat array is able to sense membrane curvature and prefers to associate with stronger convexly curved membrane surfaces

3D-modeling of Ankrd26’s ankyrin repeat array suggested a curved and rotationally twisted structure brought about by tight lateral stacking of the individual ankyrin repeats (**Figure 4A**). This suggested that Ankrd26’s ankyrin repeat array may prefer membrane surfaces, which offer better fitting curvatures, while the full N-Ank module with its ability to intercalate an amphipathic structure into one leaflet of the membrane may have the power to actively shape membranes into bent topologies, which may then also fit the curved ankyrin repeats. According to this hypothesis, the ankyrin repeats of Ankrd26 may thus be able to discriminate different curvatures of membranes. Offering small unilamellar vesicles (SUVs; average size ∼50 nm (Wolf et al., 2019)) versus large ones (LUVs) indeed led to an enhanced membrane binding of Ankrd26^Ank^ (**Figure 4B,C**). Similarly, the mouse Ankrd26 ankyrin repeat array with its very weak membrane binding also was able to sense the curvature of membranes and strongly preferred SUVs (170% above LUV binding; *P*<0.01; **Figure 4-figure supplement 1**).

**Figure 4.**
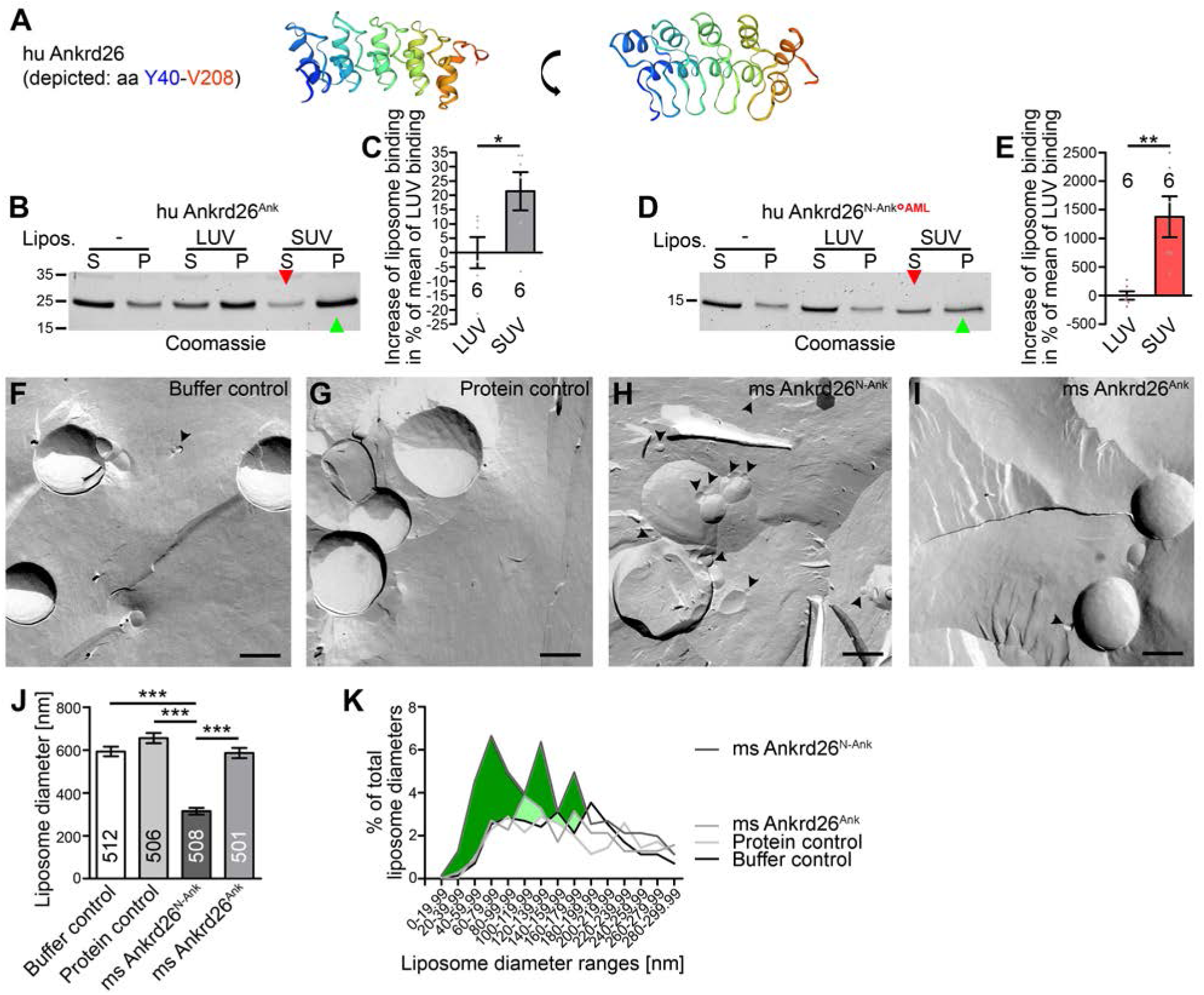
Ankrd26’s N-Ank domain recognizes membrane curvature by its ankyrin repeat arrays and shapes membrane topology. **(A)** The ankyrin repeat array of human Ankrd26 in two rotational views (modeling, https://swissmodel.expasy.org/viewer/ngl/; Y40 blue to V208 red). **(B-E)** Liposome coprecipitation studies with the wild type Ankrd26’s ankyrin repeat array **(B)** and an AML-associated mutant N-Ank module Ankrd26^N-Ank^°^AML^ **(D)**, respectively, with LUVs versus SUVs - and quantitative comparisons thereof expressed as relative increase (in percent) of binding to LUVs **(C,E)**. **(F-I)** Representative TEM images of freeze-fractured liposomes incubated with HN buffer (buffer control) **(F)**, GST as an unrelated protein control **(G)**, ms Ankrd26^N-Ank^ shaping liposomes to smaller structures (arrowheads) **(H)** or ms Ankrd26^Ank^ **(I)** and quantitative analyses of mean liposome diameters **(J)** as well as distribution analysis of liposome diameters **(K)** (data range shown, 0 to 300 nm). Bars in **F-I**, 200 nm. Colored areas in **K** mark overrepresentations of smaller liposomes upon incubation with Ankrd26^N-Ank^ when compared to all other curves (green) and to all except one of the other curves (lighter shade of green), respectively. Data in **C,E,J**, mean±SEM. Data in **K**, absolute percent numbers. (**C,E)** Bar plots with individual data points (dot plots); n=6 assays each; two-tailed, unpaired Student’s t-test. **(J)** n=512 (buffer control); n=506 (unrelated protein control); n=508 (ms Ankrd26^N-Ank^); n=501 (ms Ankrd26^Ank^) liposomes from two independent experiments and liposome preparations. Kruskal-Wallis test with Dunn’s multiple comparison test. \**P*<0.05; \*\**P*<0.01; \*\*\**P*<0.001.

The very weak membrane binding of Ankrd26^N-Ank^*°^AML^ carrying the disease mutation also was significantly enhanced upon offering strongly curved membrane surfaces represented by SUVs instead of the only very moderately curved LUVs (**Figure 4D,E**). The Ankrd26 ankyrin repeat array thus is able to sense more strongly convex membrane curvature and this ability is retained in the ankyrin repeat array of Ankrd26^N-Ank^°^AML^ which lacks one of the five ankyrin repeats when compared to wild-type Ankrd26.

### Ankrd26 is a membrane shaping protein and this function relies on the amphipathic N terminus

Using liposomes, we next analyzed whether Ankrd26 N-Ank membrane association and partial insertion leads to induction of membrane curvature (**Figure 4F-I**). Control incubations with buffer alone or with an unrelated protein (GST) mostly showed larger liposomes when analyzed by freeze-fracturing and TEM (**Figure 4F,G)**. In contrast, liposomes incubated with Ankrd26^N-Ank^ were marked by an ample presence of small liposomes (**Figure 4H-J**).

Freeze-fracturing and TEM has the advantage that liposomes with sizes over several orders of magnitudes can reliably be visualized. Quantitative determinations clearly demonstrated that Ankrd26^N-Ank^ had a strong and highly statistically significant effect on membrane topology. Ankrd26^N-Ank^-incubated liposomes with in average ∼300 nm diameter showed only about half the diameter of controls or incubations with the deletion mutant Ankrd26^Ank^ (mean, ∼600 nm) (**Figure 4J**). Thus, the complete N-Ank module has the power to actively convert membrane topologies into more strongly curved ones and the amphipathic N terminus is absolutely crucial for the membrane shaping ability of Ankrd26.

Distribution analyses of liposome diameters observed in incubations without protein, with unrelated protein, with Ankrd26^N-Ank^ and with Ankrd26^Ank^, respectively, clearly showed that especially liposomes with small diameters were much more abundant in incubations with Ankrd26^N-Ank^ than in controls. Especially in the size categories of 100 nm and beneath, their frequencies were 2-5fold as high as those of liposomes incubated with Ankrd26^Ank^ or as those of the control incubations (**Figure 4K**).

SUVs generated by sonication of LUVs roughly have diameters around 50 nm (Wolf et al., 2019). The curvatures of the more abundant small liposomes observed in EM analyses of liposomes incubated with Ankrd26^N-Ank^ thus were in the same order of magnitude as those of the SUVs that were preferred by the Ankrd26 ankyrin repeat array in the SUV versus LUV binding studies (**Figure 4B,C; Figure 4-figure supplement 1**).

### Ankrd26 plays a critical role in cellular differentiation processes

Our analyses of Ankrd26 at the plasma membrane of SK-N-SH neuroblastoma cells during retinoic acid/BDNF-induced differentiation indicated that impairment of Ankrd26’s integration into the plasma membrane is linked to AML. We thus asked whether Ankrd26 may have any role in cellular differentiation processes. Interestingly, SK-N-SH neuroblastoma cells subjected to Ankrd26 RNAi showed impairments in morphogenesis processes triggered by retinoic acid/BDNF-induced cell differentiation. Retinoic acid/BDNF treatment leads the induction and extension of cellular protrusions extending from the cell bodies (Encinas et al., 2000). Ankrd26-deficient cells, however, had less elaborate processes when compared to control cells (**Figure 5A-C**). Quantitative analyses showed that RNAi1 (Yan et al., 2020) led to a reduction of protrusion number by about a third (**Figure 5D**). Using another established RNAi site (RNAi2; Yan et al., 2020) even led to less than half of the protrusion numbers observed in control SK-N-SH neuroblastoma cells transfected with a GFP-reported plasmid expressing scrambled RNAi (**Figure 5E**). The identified Ankrd26 loss-of-function phenotype thus was consistent when two different RNAi sites were employed (**Figure 5D,E**).

**Figure 5.**
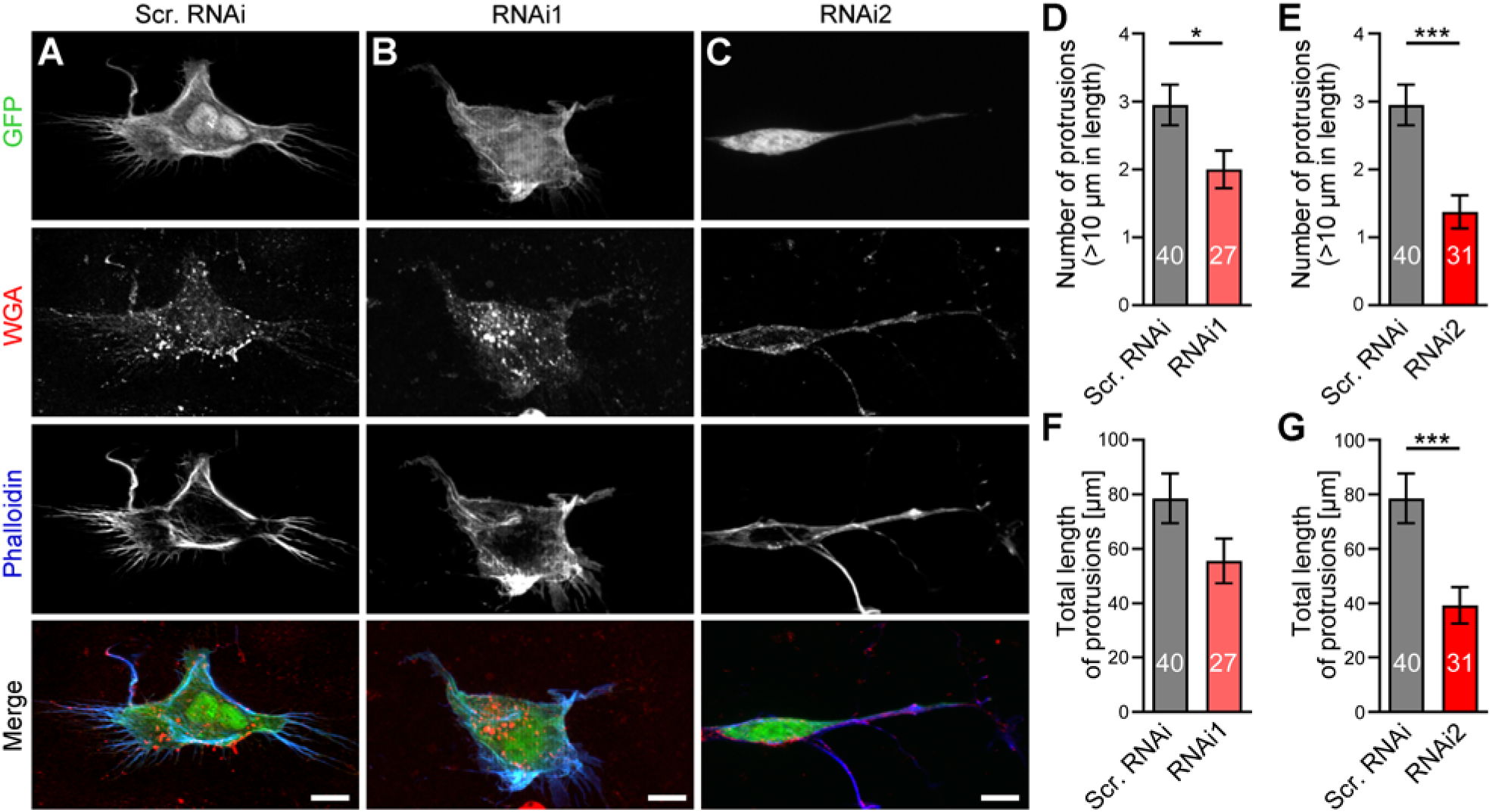
Ankrd26 is critical for retinoic acid/BDNF-induced differentiation of neuroblastoma cells. **(A-C)** MIPs of SK-N-SH neuroblastoma cells that were differentiated with retinoic acid/BDNF and transfected with GFP-reported scrambled RNAi **(A)** and Ankrd26 RNAi1 and 2 **(B,C)**, respectively. The cells were additionally stained with fluorescent wheat germ agglutinin (WGA; cell surface labeling; red in merge) and phalloidin (F-actin labeling; blue in merge). Note that Ankrd26 RNAi- transfected cells fail to adopt the morphology of differentiated cells with their protrusions. Bars, 10 µm. **D-G** Quantitative assessments of the numbers of protrusions **(D,E)** as well as of the total length of protrusions **(F,G)** of Ankrd26 RNAi1 **(D,F)** and RNAi2 **(E,G)** cells compared to control cells expressing scrambled RNAi (Scr. RNAi) (data set of phenotypical analyses repeated in RNAi2 comparisons). Data represent mean±SEM. n=27 (Ankrd26 RNAi1), n=31 (Ankrd26 RNAi2) and n=40 (Scr. RNAi) cells from 2-3 independent coverslips each per assay and two independent assays. Two- tailed, unpaired Student’s t-tests. \**P*<0.05; \*\*\**P*<0.001.

Further quantitative evaluations unveiled that also the total length of the protrusions was reduced upon Ankrd26 deficiency (**Figure 5F,G**). The reduction in total length reached about 50% using RNAi2 when compared to control (**Figure 5G**) and thereby was closely related to the reduction in protrusion number (also approx. −50%) (**Figure 5E**).

Ankrd26 deficiency thus led to strong impairments in the retinoic acid/BDNF-triggered differentiation of SK-N-SH neuroblastoma cells.

### Ankrd26 but not the AML-associated Ankrd26 mutant leads to enhanced cellular differentiation of neuroblastoma cells

We next asked whether Ankrd26 is not just critically involved in neuroblastoma cell differentiation but would also be able to effectively drive the process. Overexpression of Ankrd26 indeed promoted the extension of neuroblastoma cells (**Figure 6A-E**). In detailed quantitative examinations using IMARIS 3D-morphology tracing software parameters that ensured that also smaller effects should be covered, we observed that the numbers of protrusions determined as numbers of terminal points increased by more than 40% when compared to GFP controls (**Figure 6D**). Additionally, an excess of Ankrd26 led to a strong increase of the overall extension of IMARIS traces (“filament”) representing the cellular morphologies. With an increase of about 60%, also this parameter of cellular morphology was pronounced and statistically significant (**Figure 6E**).

**Figure 6.**
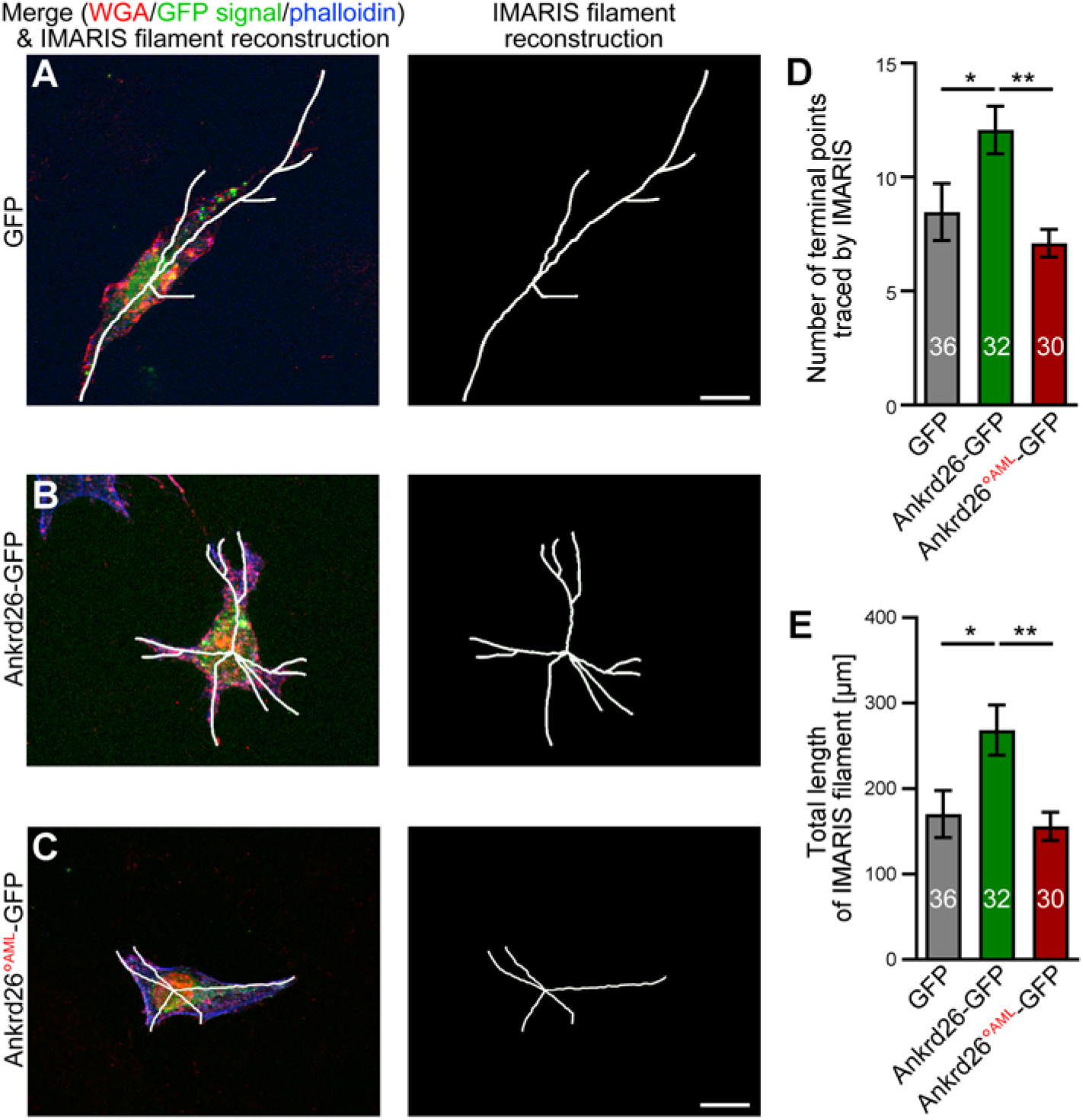
Ankrd26 but not the AML-associated Ankrd26 mutant leads to enhanced cellular differentiation of neuroblastoma cells. **(A-C)** MIPs of retinoic acid/BDNF-incubated SK-N-SH neuroblastoma cells that were transfected with GFP **(A),** Ankrd26-GFP **(B)** and Ankrd26°^AML^-GFP **(C)**, respectively. The cells were additionally stained with fluorescent wheat germ agglutinin (WGA; cell surface labeling; red in merge) and phalloidin (F-actin labeling; blue in merge). Additionally, representations of the IMARIS software-based 3D-reconstructions of the morphology of cells (IMARIS filaments) are shown. Bars, 10 µm. **(D,E)** Quantitative assessments of the numbers of protrusions determined by the number of terminal points **(D)** as well as of the total length of the IMARIS filament **(E)**. Data represent mean±SEM. n=36 (GFP), n=32 (Ankrd26-GFP) and n=30 (Ankrd26°^AML^-GFP) cells from 2-3 independent coverslips each per assay and four independent assays. One-way ANOVA with Tukey post hoc test. \**P*<0.05; \*\**P*<0.01.

Interestingly, the AML-associated Ankrd26 mutant, which we demonstrated to be membrane-association-deficient, was unable to bring about Ankrd26-mediated cellular effects during retinoic acid/BDNF-induced differentiation of neuroblastoma cells. In contrast to WT Ankrd26, Ankrd26°^AML^ expression failed to lead to any additional protrusions; this difference was highly statistically significant (**Figure 6A,C,D**). In addition, also the total length of the IMARIS filament remained at the level of GFP-transfected control cells (**Figure 6A,C-E**). Thus, in contrast to WT Ankrd26, the AML-associated Ankrd26 mutant completely failed to boost neuroblastoma cell differentiation.

### The N terminus missing in the AML-associated Ankrd26 mutant and the coiled coil domain of Ankrd26 are both critical for differentiation of neuroblastoma cells

In order to address, whether the AML-associated mutant of Ankrd26 is unable to maintain Ankrd26 functions in cell differentiation, and to reveal, whether beyond a functional N-Ank module also the coiled coil domain located in the C terminal half of Ankrd26 is required for Ankrd26 functions, we next conducted rescue attempts of the Ankrd26 loss-of-function phenotype (**Figure 7A-G**). Both the reduction of the number of protrusion and also a reduction in the total length of the 3D-IMARIS-based cell morphology reconstruction (IMARIS filament) caused by Ankrd26 RNAi were fully rescued by coexpression of an RNAi-insensitive Ankrd26 (Ankrd26*-GFP) (**Figure 7A-C,F,G**). This successful rescue by Ankrd26*-GFP firmly demonstrated that the identified Ankrd26 RNAi phenotypes are specifically caused by a lack of Ankrd26 (**Figure 7A-C,F,G**).

**Figure 7.**
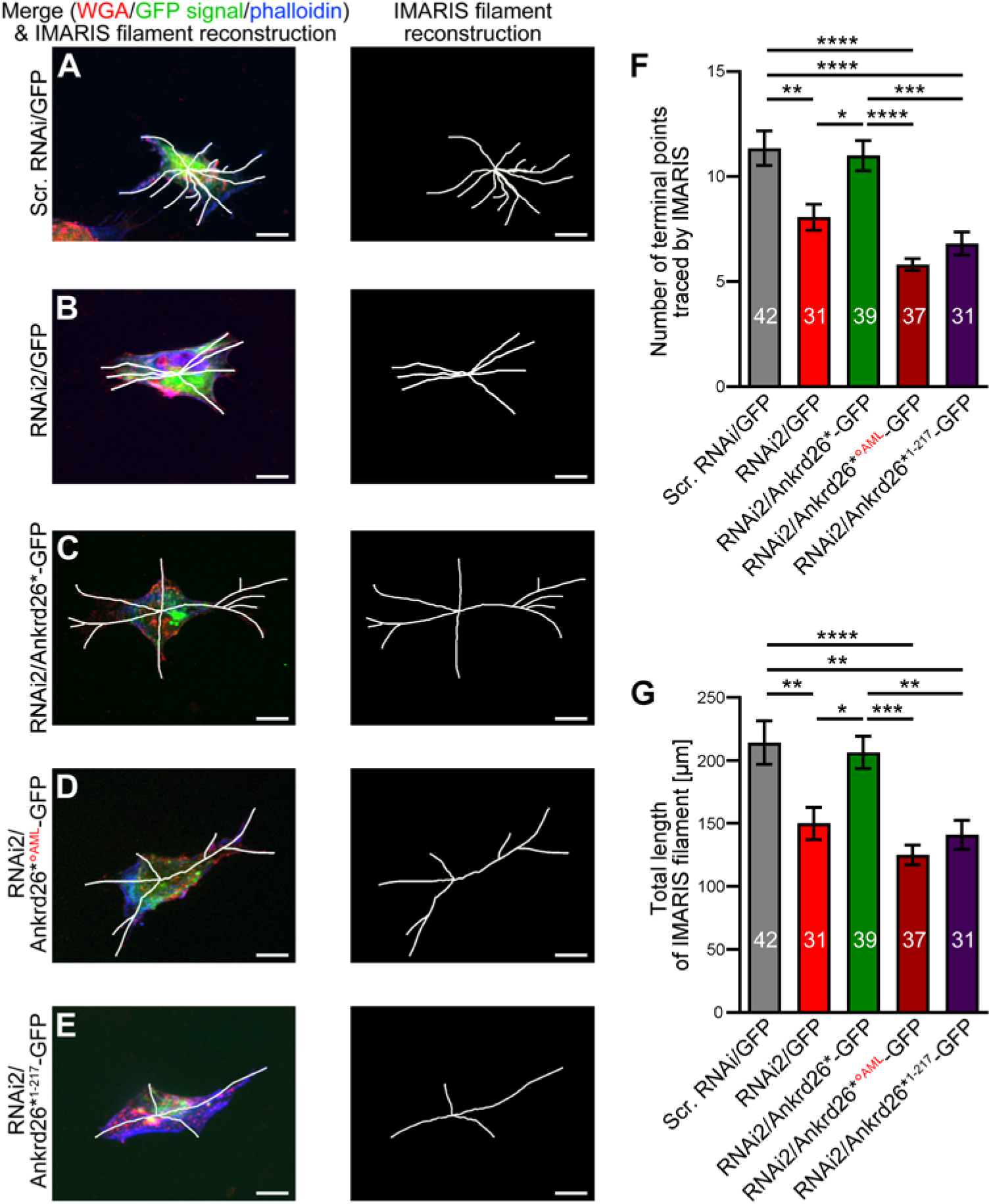
The N terminus missing in the AML-associated Ankrd26 mutant and the coiled coil domain of Ankrd26 are both critical for differentiation of neuroblastoma cells. **(A-E)** MIPs of SK-N-SH neuroblastoma cells differentiated with retinoic acid/BDNF and transfected with GFP-reported scrambled RNAi (Scr. RNAi) **(A),** with GFP-reported Ankrd26 RNAi2 (RNAi2) **(B)**, Ankrd26 RNAi2 coexpressing a RNAi-insensitive (silent mutations) Ankrd26 (Ankrd26*) (RNAi2/Ankrd26*-GFP; WT rescue) **(C)** and two RNAi-insensitive mutant Ankrd26 proteins, respectively (**D**; RNAi2/Ankrd26*°^AML^-GFP; **E**; RNAi2/Ankrd26*^1-217^-GFP, i.e. a deletion mutant lacking the coiled coil domain-containing C terminal part of Ankrd26). The cells were additionally stained with fluorescent wheat germ agglutinin (WGA; cell surface labeling; red in merge) and phalloidin (F-actin labeling; blue in merge). Note that Ankrd26 RNAi-transfected cells fail to adopt the morphology of differentiated cells with their protrusions and that this defect was rescued by replenishing the cells with Ankrd26* but not with the two mutants analyzed. Bars, 10 µm. **(F-G)** Quantitative assessments of the numbers of protrusions **(F)** as well as of the total length of the IMARIS filament **(G)**. Data represent mean±SEM. n=42 (Scr. RNAi), n=31 (RNAi2), n=39 (RNAi2/Ankrd26*-GFP), n=37 (RNAi2/Ankrd26*°^AML^-GFP) and n=31 (RNAi2/Ankrd26*^N-Ank^-GFP) cells from 2-3 independent coverslips each per assay and four independent loss-of-function/rescue assays. One-way ANOVA with Tukey post hoc test. \**P*<0.05; \*\**P*<0.01; \*\*\**P*<0.001; \*\*\*\**P*<0.0001.

In contrast, the AML-associated mutant of Ankrd26 completely failed to restore the WT situation of retinoic acid/BDNF-induced neuroblastoma cell differentiation. The strongly reduced number of protrusions in Ankrd26*°^AML^-GFP-reexpressing cells was highly significantly different from both, control cells and RNAi2/Ankrd26*-GFP rescue cells. When compared to Ankrd26 RNAi, no positive effects of reexpression of Ankrd26*°^AML^-GFP could be observed (**Figure 7B-D,F**). A similar failure of the AML-associated Ankrd26 mutant to rescue Ankrd26 RNAi phenotypes was observed for the second cell morphology parameter. Also the overall length of the 3D morphology reconstructions was not rescued by Ankrd26*°^AML^-GFP.

Intriguingly, also an Ankrd26 deletion mutant that comprised an intact N-Ank domain but lacked the entire more C terminal part of Ankrd26 including the coiled coil domain resulted in a complete failure of rescuing the Ankrd26 loss-of-function phenotypes (**Figure 7E-G**).

Therefore, the thus far studied, critical N-Ank functions need to be combined with yet to be revealed functions of the coiled coil domain-containing C terminal part of Ankrd26 in order to bring about Ankrd26’s critical role in cell differentiation.

### The papillary thyroid carcinoma-associated fusion of parts of Ankrd26 and RET exhibits a membrane localization mediated by the Ankrd26 portion

Alterations in Ankrd26’s plasma membrane association and/or defects in Ankrd26’s yet largely uncharacterized coiled coil domain functions may also play a role in the pathophysiology of the Ankrd26^1-1405^-RET^712-1114^ fusion mutant found in papillary thyroid carcinoma. This Ankrd-RET fusion comprises a C terminally truncated version of Ankrd26 (lacking parts of Ankrd26’s coiled coil domain) together with the kinase domain of RET. RET is normally anchored in the membrane by a transmembrane domain and RET kinase signaling thus originates from the plasma membrane. The entire N terminal half of RET including its transmembrane domain, however, is lacking in the piece of RET included in the Ankrd26-RET fusion found in papillary thyroid carcinoma. Instead, only the C terminal tyrosine kinase domain of RET was present in the Ankrd26-RET fusion (Straubitz et al., 2019). We therefore first addressed the membrane binding of RET and RET^712-1114^ (**Figure 8A-D; Figure 8-figure supplement 1**). Wild-type RET-GFP was found along the secretory pathway (ER, Golgi) – as expected for a transmembrane protein – and showed a good spatial overlap with CherryF at the plasma membrane (**Figure 8A**). Cofractionation with CherryF (**Figure 8B**) clearly confirmed the plasma membrane association of RET-GFP. In contrast, RET^712-1114^-GFP did neither localize with CherryF at the plasma membrane nor cofractionated with this plasma membrane marker (**Figure 8C,D**).

**Figure 8.**
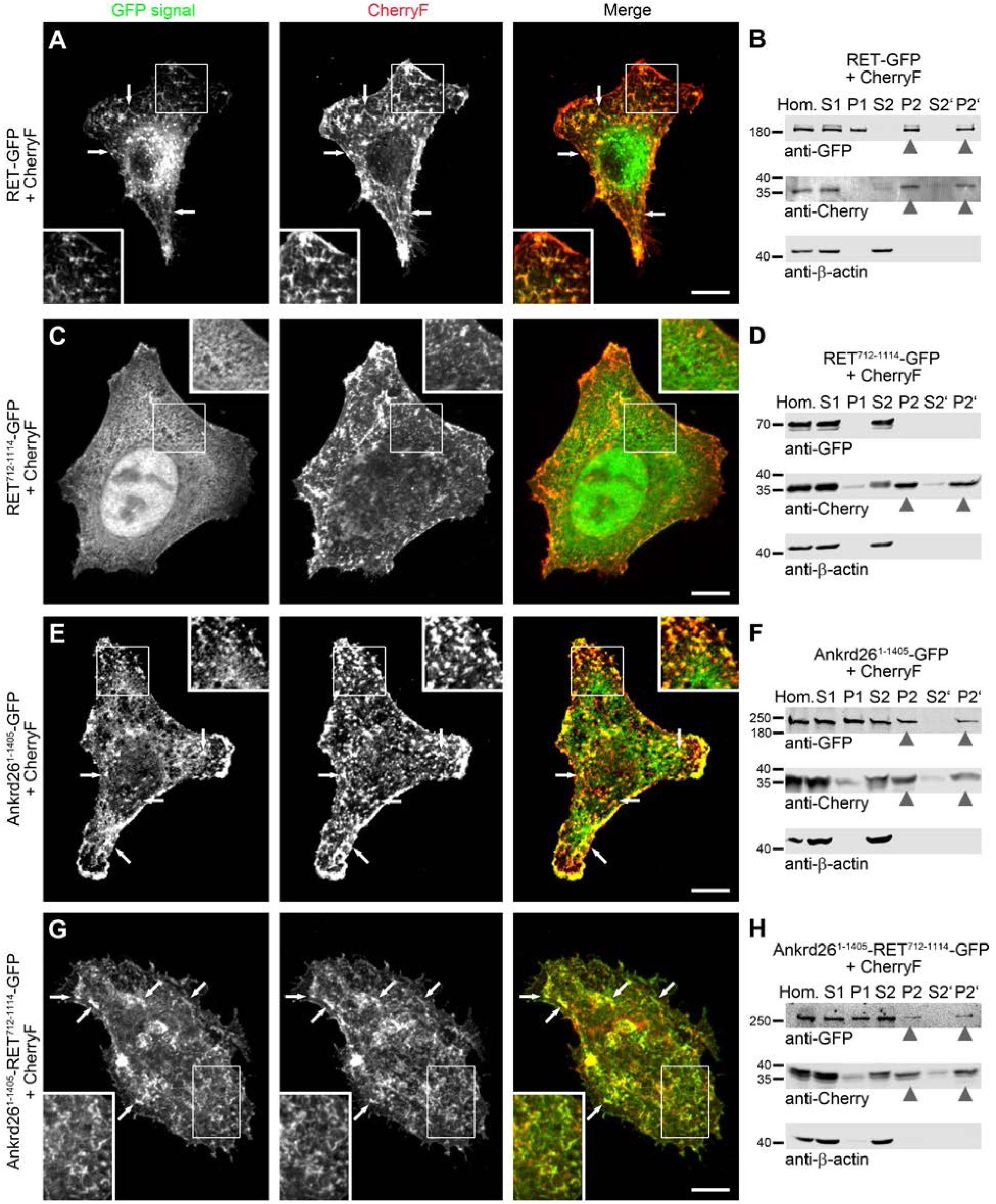
The papillary thyroid carcinoma-associated fusion of Ankrd26 and RET exhibits a membrane localization mediated by the included Ankrd26 portion. **(A-H)** MIPs of indicated proteins in CherryF-cotransfected HeLa cells **(A,C,E,G)** and subcellular fractionations of lysates of HEK293 cells expressing the indicated proteins **(B,D,F,H)**. Analyzed were RET-GFP (full-length; aa1-1114) **(A,B)**, the RET fragment RET^712-1114^-GFP **(C,D)**, the Ankrd26 fragment Ankrd26^1-1405^-GFP **(E,F)** and the papillary thyroid carcinoma-associated fusion Ankrd26^1-1405^-RET^712-1114^-GFP **(G,H)**. Arrows in **A, E** and **G** mark some examples of colocalization of RET- GFP **(A)**, Ankrd26^1-1405^-GFP **(E)** and Ankrd26^1-1405^-RET^712-1114^-GFP **(F)**, respectively, with the plasma membrane marker CherryF. Bars, 10 µm. Insets are magnifications of boxed areas highlighting colocalizations with the membrane marker CherryF **(A,E,G)**. For additional fluorescence intensity plots for the red and the green fluorescence channel highlighting colocalizations of RET- GFP, Ankrd26^1-1405^-GFP and Ankrd26^1-1405^-RET^712-1114^-GFP but not RET^712-1114^-GFP with the plasma membrane marker CherryF see **Figure 8-figure supplement 1**. Proteins detected in the immunoblotted membrane fractions P2 and P2’ are marked by arrowhead **(B,D,F,H)**.

RET signaling normally originates from the plasma membrane. Interestingly, fusion of Ankrd26^1-1405^ with RET^712-1114^ restored a plasma membrane-targeting of the RET kinase domain, as the Ankrd fragment Ankrd26^1-1405^ turned out to be fully membrane binding-competent and by fusion conferred this ability to RET^712-1114^, as evidenced by colocalization and cofractionation of the papillary thyroid carcinoma-associated Ankrd26-RET fusion with CherryF (**Figure 8E-H; Figure 8-figure supplement 1**).

Thus, in papillary thyroid carcinoma expressing Ankrd26-RET, aberrant signaling pathways seem not to originate from the fact that Ankrd26 was not correctly anchored at the plasma membrane or that RET was not localized to the plasma membrane. Rather, it seemed that some other mechanisms are involved in the aberrant signaling of the Ankrd26-RET fusion leading to papillary thyroid cancer.

### Self-association of RET is restored by a related property of Ankrd26 in the Ankrd26-RET fusion found in papillary thyroid carcinoma

The fusion of the protooncogene *RET* with *ANKRD26* found in papillary thyroid carcinoma patients only comprises the sequence encoding for the RET amino acids 712 to 1114 (Staubitz et al., 2019), which - quite common in *RET* fusions identified in cancer (Shaw et al., 2013; Liu et al., 2021) - encodes for the kinase domain of RET (Takahashi and Cooper, 1987) (**Figure 9A,B**). The lacking N terminal half of RET encodes for its extracellular part and the transmembrane domain. Membrane association is normally ensured by the transmembrane domain of RET and this aspect of RET signaling was restored by fusing the C terminal RET fragment to Ankrd26^1-1405^ (**Figure 8**), which comprises a fragment of the predicted extended coiled coil domain of Ankrd26 and the membrane-binding N-Ank module **(Figure 9C,D**). Another aspect important for RET signaling is the assembly of RET in multireceptor complexes (Mulligan, 2014). Crosslink studies with RET-GFP and the zero-length crosslinker EDC demonstrated this RET property in HEK293 cells (**Figure 9E**). EDC crosslink led to a high molecular weight band at above 460 kD reflecting RET self-association products. These RET self-association products increased with rising crosslinker concentrations and were to some extent even visible without crosslinker suggesting that they are in part even SDS-resistant (**Figure 9E**).

**Figure 9.**
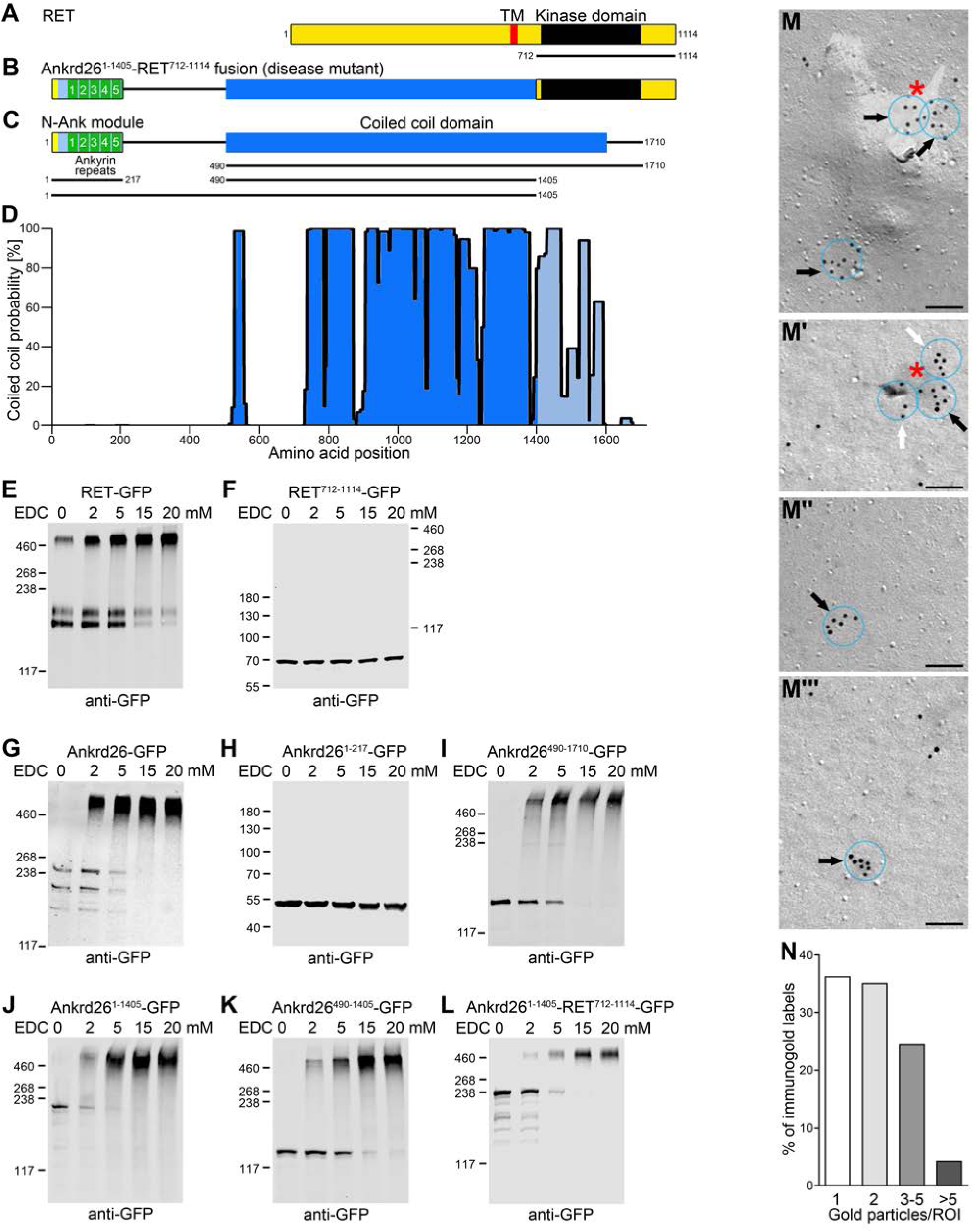
Ankrd26 is able to self-associate, the Ankrd26 fragment included in the Ankrd26^1-1405^-RET^712-1114^ fusion mutant confers this function to the fusion mutant thereby mimicking RET dimerization and this self-association capability of Ankrd26 is reflected by Ankrd26 clusters at the plasma membrane. **(A,B)** Schematic representations of RET **(A)** and the papillary thyroid carcinoma-associated Ankrd26- RET fusion **(B)**. TM, transmembrane domain. **(C)** Schematic representation of human Ankrd26 with its N terminal N-Ank module and C terminal coiled coil domain. Black lines represent additional deletion mutants tested for self-association. **(D)** Coiled coil prediction by COILS (blue). The coiled coil domain part lacking in the Ankrd26-RET mutant is in light blue. **(E-L)** Immunoblot analyses of experiments with increasing concentrations of the crosslinker EDC added to cell lysates containing overexpressed RET-GFP **(E)**, the C terminal RET fragment found in papillary thyroid carcinoma (RET^712-1114^-GFP) **(F)**, full-length Ankrd26-GFP (**G**), Ankrd26^1-217^-GFP encompassing Ankrd26’s N- Ank module **(H)**, the complete coiled coil domain (Ankrd26^490-1710^-GFP) **(I)**, the part of Ankrd26 up to the fusion point in Ankrd26-RET (Ankrd26^1-1405^-GFP) **(J)** and the part of only Ankrd26’s coiled coil domain included in the papillary thyroid carcinoma-associated Ankrd26-RET mutant **(K)**. Note the EDC-induced high-molecular weight bands except for the N-Ank module **(H)** and the RET kinase domain **(F)**. **(L)** Crosslink studies with the papillary thyroid carcinoma-associated Ankrd26^1-1405^- RET^712-1114^ fusion mutant showing that fusion to Ankrd26^1-1405^ efficiently brings about self- association. **(M-M’’’)** Examples of transmission electron microscopical detections of Ankrd26 at platinum-shadowed, freeze-fractured plasma membranes of retinoic acid/BDNF-stimulated SK-N-SH neuroblastoma cells in form of single and double labels (not marked) as well as of clustered anti- Ankrd26 immunogold labels (arrows) (ROI, transparent blue circles, diameter, 100 nm). Bar, 100 nm. White arrows, clusters of 3-5 gold labels/ROI; black arrows, clusters of more than 5 gold labels; red asterisks, Ankrd26 superclusters with overall diameters >100 nm (separated into several ROIs, i.e. subclusters, in quantitative cluster analyses). **(N)** Relative distribution (in percent) of all anti-Ankrd26 immunogold labels imaged with respect to clustering (single, double, 3-5, >5 gold particles/ROI). Absolute numbers of labels per group, single (1), n=1091; double (2), n=1056; 3-5, n=739; >5, n=127 from 94 images from 2 independent experiments.

In contrast, the RET fragment (RET^712-1114^), which is fused to Ankrd26 in the papillary thyroid carcinoma-linked Ankrd26-RET fusion, was unable to self-associate. Only the monomeric band of RET^712-1114^-GFP was seen irrespective of the concentration of crosslinker used (**Figure 9F**). The RET fragment remaining in the papillary thyroid carcinoma Ankrd-RET fusion thus lacks the RET self-association ability critical for RET signaling.

Interestingly, related experiments with Ankrd26-GFP clearly demonstrated that similar to RET also Ankrd26 gave rise to high molecular weight self-association products upon incubations with EDC analyzed by immunoblotting analyses (**Figure 9G**). Corresponding to the increasing high molecular weight self-association products, the band reflecting monomeric full-length Ankrd26-GFP (about 240 kD) and also the bands reflecting GFP-containing proteolytic fragments of Ankrd26 declined with rising EDC concentrations (**Figure 9G**).

The use of deletion mutants demonstrated that the self-association capability of Ankrd26 we observed was not mediated by Ankrd26’s N-Ank module (**Figure 9H**). Instead, it clearly was a property of the C terminal coiled coil domain of Ankrd26 (**Figure 9I**). Instead of dimers, it mostly were higher order assemblies with apparent molecular weights above 500 kD that were formed (**Figure 9I**).

The fusion at the position of aa1405 of Ankrd26 with RET found in papillary thyroid carcinoma represents a disruption of Ankrd26’s coiled coil domain, which spans the region between aa500 and aa1600 (**Figure 9C,D**). Interestingly, both Ankrd26^1-1405^ as well as Ankrd26^490-1405^ still were able to efficiently self-associate despite the deletion of all coiled coil domain residues beyond the RET fusion point (**Figure 9J,K**). This suggested that, despite the partial disruption of Ankrd26’s coiled coil domain, the remaining part of Ankrd26 still is able to self-associate and could therefore also confer such functions to the papillary thyroid carcinoma-linked Ankrd26/RET fusion. Studies with Ankrd26^1-1405^-RET^712-1114^ indeed showed that the fusion product was able to self-associate. Self-association products at a size of about 500 kD were clearly detectable and rose with rising EDC concentration, while low molecular weight bands of the Ankrd26^1-1405^-RET^712-1114^ fusion declined correspondingly (**Figure 9L**).

### Ankrd26 forms nanoclusters at the plasma membrane

Self-association could be an important aspect in Ankrd26 functions. Freeze-fractured plasma membranes of retinoic acid/BDNF-treated SK-N-SH neuroblastoma cells indeed showed a frequent occurrence of multiple anti-Ankrd26 immunogold labels in close proximities of 5-40 nm to each other (**Figure 9M-M’’’**).

Quantitative analyses of in total 1388.9 µm^2^ of P-faces (i.e. cyotosolic faces) showed that about a third of all anti-Ankrd26 immunogold labels occurred in form of single gold particles, another third in form of doublets and the last third in form of clusters of three or more gold particles (**Figure 9N**). Some of the Ankrd26 clusters were up to ∼200 nm in extension (for examples see **Figure 9M** and **Figure 9M’**). Superclusters contained up to 17 gold particles. The maximal labeling frequency of 100 nm ROI clusters was 10 gold particles. The Anrd26 molecules detected in form of nanoclusters thus were in intimate contact with each other.

### The Ankrd26^1-1405^-RET^712-1114^ fusion found in papillary thyroid carcinoma leads to increased ERK1/2 activation

Our studies so far demonstrated that both key aspects of RET signaling, the self-association and the membrane localization of RET are disrupted in RET^712-1114^ but fully restored by fusion to Ankrd26^1-1405^. In fact, determinations of the levels of phosphorylated ERK1/2 demonstrated that the ERK1/2 pathway was neither activated in RET kinase domain-expressing cells nor in Ankrd26^1-1405^-expressing cells. Also overexpression of full-length Ankrd26 did not cause any obvious activation of the ERK pathway but pERK1/2 levels were very low (**Figure 10A-D**).

**Figure 10.**
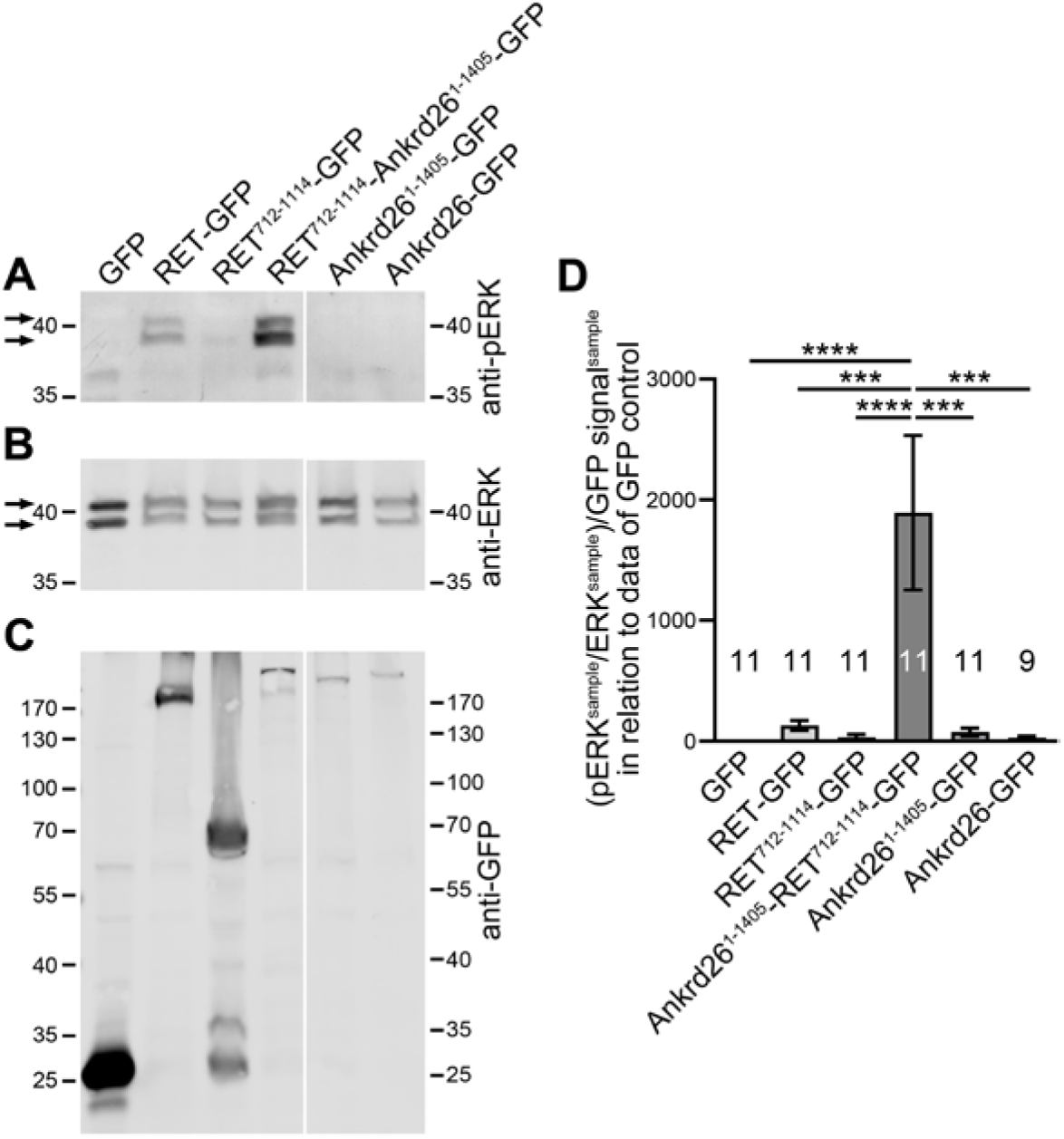
The papillary thyroid carcinoma-associated fusion of Ankrd26 and RET leads to strong activation of ERK1/2. **(A-C)** Western blot analyses of phosphoERK (pERK) **(A)** and ERK levels **(B)** as well as of the GFP immunosignals **(C)** in lysates of HEK293 cells transfected with GFP, RET-GFP, RET^712-1114^-GFP (RET kinase domain fragment) and GFP fusion proteins of the papillary thyroid carcinoma-associated Ankrd26^1-1405^-RET^712-1114^ mutant as well as of the Ankrd26^1-1405^ fragment and of full-length Ankrd26. **(D)** Quantitative immunoblotting determination of pERK/ERK activity normalized to the expression levels of the GFP fusion proteins, as determined by anti-GFP immunoblotting, and expressed in relation to the data for the GFP control set to 1. Data represent mean±SEM. n=11 and 9 assays each, respectively, as indicated in the figure. One-way ANOVA and Tukey post hoc test. \*\*\**P*<0.001; \*\*\*\**P*<0.0001.

Fusion of the two fragments from RET and Ankrd26, however, led to pERK/ERK levels that by far exceeded those of full-length RET overexpression (**Figure 10A-D**). Normalized to GFP control, RET-GFP overexpression led to an about 100fold activation of the ERK1/2 pathway when compared to control. Overexpression of the papillary thyroid carcinoma-associated Ankrd26^1-1405^-RET^712-1114^ mutant led to phosphoERK1/2 levels that were almost 2000fold above control. This increase of ERK1/2 activation caused by the Ankrd26^1-1405^-RET^712-1114^ fusion mutant was highly statistically significant when compared to all other conditions tested (**Figure 10D**).

The Ankrd26^1-1405^-RET^712-1114^ fusion found in papillary thyroid carcinoma patients thus did not seem to reflect dominant-negative RET functions. Instead, fusion of RET^712-1114^ with Ankrd26^1-1405^ caused a strong activation of the ERK pathway. It hereby was specifically the fusion, but not the involved Ankrd26 fragment by itself, which caused the massive increase in ERK/MAPK signaling.

### The papillary thyroid carcinoma-associated Ankrd26^1-1405^-RET^712-1114^ fusion shows increased RET autophosporylation

A molecular mechanism underlying the observed activation of ERK1/2 signaling could be the generation of a constitutively active form of RET, which would be decoupled from ligand binding due to the deletion of the RET N terminus. GFL-binding of one of the four GDNF receptor-α (GFRα) family members serving as coreceptors leads to RET recruitment, RET dimerization and subsequent RET autophosphorylation at multiple tyrosines including e.g. Y753, Y905, Y981, Y1015 and Y1062. RET autophosphorylation then leads to a recruitment of a variety of signaling components responsive to phosphorylated tyrosines and to the assembly of larger signaling complexes (Mulligan, 2014). RET phosphorylations thereby reliably reflects the activity of RET signaling. Quantitative (fluorescence-based) determination of anti-RET pY905 and anti-RET pY1015 immunoblotting signals in relation to anti-RET and anti-GFP immunoblotting thus allowed us to directly address and compare the levels of autophosphorylation of RET, of the RET^712-1114^ fragment and of the Ankrd26^1-1405^-RET^712-1114^ fusion found in papillary thyroid cancer and to thereby compare the RET kinase signaling activities elicited by these three proteins. Importantly, lysates obtained from cells transfected with the papillary thyroid carcinoma fusion mutant Ankrd26^1-1405^-RET^712-1114^ demonstrated that, despite the low expression of the mutant consistently shown by both anti-GFP and anti-RET antibodies, fusing the membrane targeting- and self-association-competent Ankrd26 part with the kinase domain of RET led to strong RET signaling, as determined by Y1015 phosphorylation (**Figure 11A-C**). Related results were obtained with an anti-RET antibody directed against phosphorylated Y905 (pY095) (**Figure 11-figure supplement 1**).

**Figure 11.**
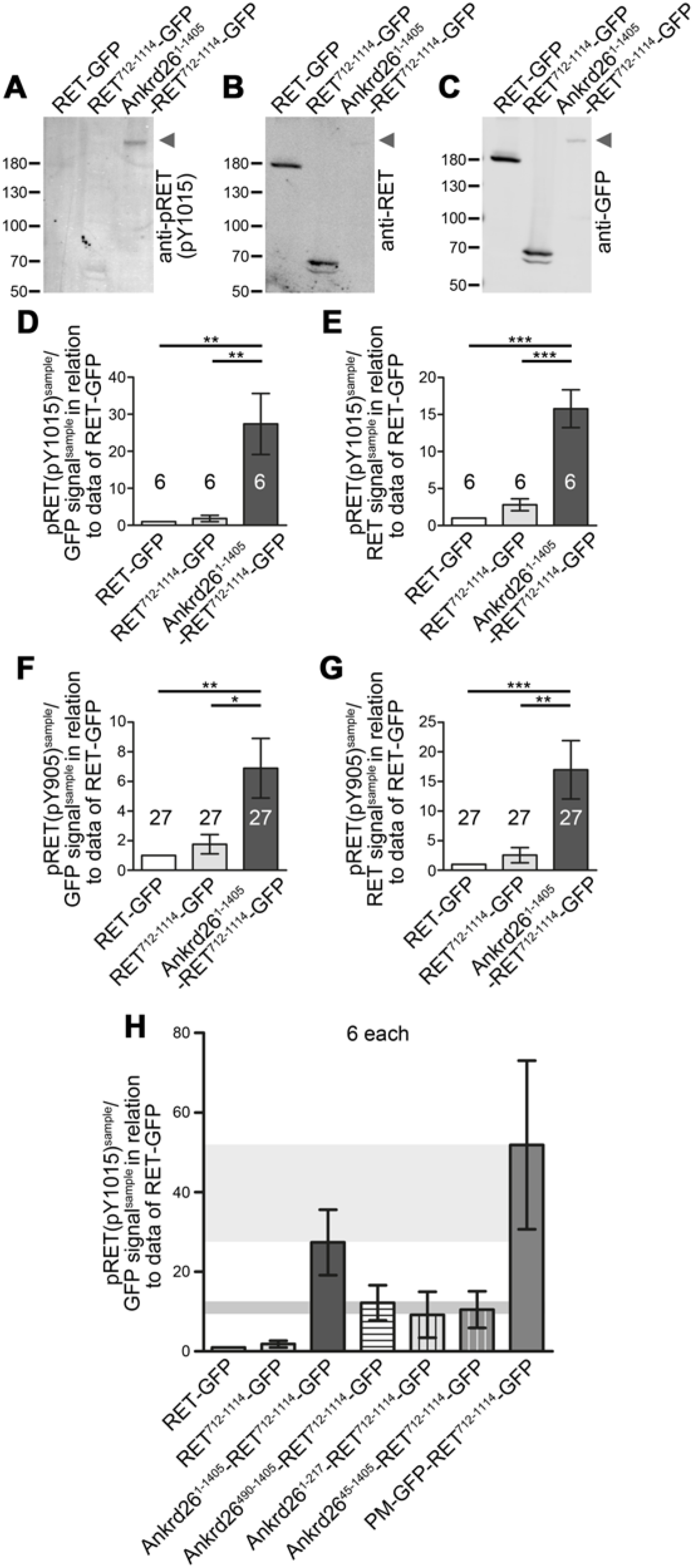
Ankrd26^1-1405^-RET^712-1114^ fusion leads to strongly increased autophosphorylation of the RET kinase domain – an effect involving both N-Ank and coiled coil domain functions. **(A-C)** Representative images of lysates of HEK293 cells transfected with GFP-tagged full-length RET (RET^1-1114^), RET^712-1114^ and the papillary thyroid carcinoma-associated fusion Ankrd26^1-1405^- RET^712-1114^, respectively, that were immunoblotted with antibodies specifically detecting pRET (pY1015) **(A)**, with antibodies against the C terminal domain of RET **(B)** and with anti-GFP antibodies **(C)**, respectively. Arrowheads mark the positions of the highly Y1015 phosphorylated Ankrd26^1-1405^-RET^712-1114^-GFP. **(D,E)** Relative quantitative determination of RET activity by anti- Y1015 immunoblotting normalized to expression, as determined by anti-GFP immunoblotting **(D)** and anti-RET immunoblotting **(E)**, respectively. For comparability, anti-pRET/anti-GFP and anti- pRET/anti-RET ratios of GFP-tagged RET-GFP, RET^712-1114^ and the papillary thyroid carcinoma- associated Ankrd26^1-1405^-RET^712-1114^ fusion mutant were normalized to the average of the of full- length RET-GFP data obtained from each assay. Note that the papillary thyroid carcinoma-associated Ankrd26^1-1405^-RET^712-1114^-GFP mutant showed a much higher RET Y1015 autophosphorylation than RET or the RET kinase domain alone **(D,E)**. **(F,G)** Related quantitative immunoblotting analyses of phosphorylated RET Y905 (see **Figure 11-figure supplement 1** for examples of blot images). **(H)** Quantitative immunoblotting analyses of RET Y1015 autophosphorylation/GFP levels of 3 different Ankrd26 mutants and PM-GFP (palmitoylated GFP) fused to RET^712-1114^ in comparison to the data obtained for GFP-tagged RET-GFP, RET^712-1114^ and Ankrd26^1-1405^-RET^712-1114^ (data from **D** repeated for comparison). Note that lack of the N-Ank-containing N terminus of Ankrd26 (Ankrd26^490-1405^), lack of Ankrd26’s self-association capability (Ankrd26^1-217^) and lack of the N terminal amphipathic helix of Ankrd26 (Ankrd26^45-1405^) each led to strongly reduced RET Y1015 autophosphorylation levels when compared to the disease mutant, whereas PM-GFP fusion led to strong RET Y1015 autophosphorylation. Data represent mean±SEM. n=6 assays each **(D,E,H)** and n=27 assays each **(F,G)**, respectively. Bartlett’s test and Tukey post hoc test. \**P*<0.05; ** *P*<0.01; \*\*\**P*<0.001.

Quantitative determinations of the pY1015 phosphorylation in RET, in RET^712-1114^and in Ankrd26^1-1405^-RET^712-1114^ in relation to the anti-GFP-detected protein expression levels showed that RET kinase activity levels of the disease mutant Ankrd26^1-1405^-RET^712-1114^ were 15-27fold higher than those of RET or the RET kinase domain alone (**Figure 11D**). Similar differences were obtained when pRET data were expressed as pRET/RET signals. Again, the Ankrd26^1-1405^-RET^712-1114^ fusion mutant showed autophosphorylation levels that were more than 5-15times higher than those of RET or the RET kinase domain. Statistical analyses demonstrated that, in both quantitative determinations, the increase in pY1015 levels was highly significant when compared to RET-GFP or to RET^712-1114^-GFP (**Figure 11E**).

Similar results were obtained when Y905 phosphorylation was determined quantitatively. Also this RET kinase autophosphorylation site showed strongly increased activity in the papillary thyroid carcinoma mutant Ankrd26^1-1405^-RET^712-1114^ (**Figure 11F,G**). Fusion of the membrane-binding and self-association-competent part of Ankrd26 with the kinase domain of RET, as found in papillary thyroid carcinoma patients (Staubitz et al., 2019), thus leads to a fusion protein with a massively increased autophosphorylation of RET.

Interestingly, dissection of the Ankrd26 properties involved in the strongly increased autophosphorylation of the papillary thyroid carcinoma mutant Ankrd26^1-1405^-RET^712-1114^ by the use of mutants lacking the coiled coil domain-mediated Ankrd26 self-association ability (Ankrd26^1-217^-RET^712-1114^) or lacking the N terminal part including the N-Ank module (Ankrd26^490-1405^-RET^712-1114^) both still led to some moderately elevated Y1015 phosphorylation but the Y1015-autophosphorylation levels of both mutants were less than half of those found in the papillary thyroid carcinoma mutant Ankrd26^1-1405^-RET^712-1114^ (**Figure 11H**). Interestingly, deletion of the amphipathic helix was sufficient for the reduction of autophosphorylation levels (**Figure 11H**).

This suggested that i) membrane binding and ii) self-association both together promote RET kinase domain autophosphorylation in Ankrd26^1-1405^-RET^712-1114^. It seems likely that these Ankrd26 properties relate to the Ankrd26-enriched membrane nanodomains we observed at the plasma membrane (**Figure 9M,N**). To vigorously address this conclusion, we constructed a palmitoylated GFP and fused it to the RET kinase domain. Without relying on self-association as a molecular mechanism, palmitoylation should ensure both efficient plasma membrane anchoring and nanodomain formation by its preference for lipid rafts (Arni et al., 1998). Indeed, the Y1015 phosphorylation levels of PM-GFP-RET^712-1114^ were strongly elevated suggesting that the molecular properties at work in the papillary thyroid carcinoma Ankrd26^1-1405^-RET^712-1114^ were successfully mimicked (**Figure 11H**). These results may relate to RET activation achieved by a combination of artificially added dimerization and myristoylation moieties to the RET kinase domain (Richardson et al., 2009) and suggest that indeed efficient membrane targeting and the formation of RET kinase domain-enriched membrane nanodomains by the membrane-binding and self-associating part of Ankrd26 are the molecular mechanisms underlying the pathophysiological signaling found in papillary thyroid carcinoma expressing Ankrd26^1-1405^-RET^712-1114^.

## Discussion

Derailed signaling pathways associated with many types of cancer often originate from the plasma membrane. The development of causal therapies relies on knowledge of the involved pathomechanisms of the different types of cancer. During the last years, a variety of *ANKRD26* mutations have been described as linked to different malignancies (Cerami et al., 2012; Marconi et al., 2017; Staubitz et al., 2019). In line with some crucial importance of the N terminus of Ankrd26 for proper function, monoallelic single nucleotide substitutions in the 5’UTR of the *ANKRD26* gene leading to the use of another start codon further 3’ and thereby to N terminal truncation of Ankrd26 are linked to thrombocytopenia – an autosomal dominant bleeding disorder – and AML (Pippucci et al., 2011; Noris et al., 2011; Bluteau et al., 2014; Marconi et al., 2017). Fusion of a C terminally truncated Ankrd26 with RET also leads to cancer (Straubitz et al., 2019). Yet, little was known about the properties of the Ankrd26 protein as such. Our analyses unveil several molecular Ankrd26 functions affected by the disease mutations and thereby shed light on Ankrd26-related pathomechanisms. We furthermore show that Ankrd26 is critical for cellular differentiation processes and our detailed phenotypical analyses revealed that particularly the induction of protrusions is affected by Ankrd26 deficiency. Ankrd26 overexpression consistently led to the opposite phenotype, i.e. to a surplus of protrusions during retinoic acid/BDNF-induced cellular differentiation of neuroblastoma cells.

Our biochemical and cell biological examinations unveiled that Ankrd26 is a plasma membrane-binding protein, whose ankyrin repeats are part of a functional N-Ank module for membrane binding as well as for membrane curvature induction and sensing. The critical physiological relevance of N- Ank module-mediated functions of Ankrd26 is reflected by the finding that the N-Ank domain was found to be dysfunctional in the analyzed AML-associated Ankrd26 mutant protein. In line, the Ankrd26°^AML^ mutant was neither able to elicit the gain-of-function phenotypes we identified for WT Ankrd26 nor was the Ankrd26*°^AML^ mutant able to rescue any of the identified Ankrd26 loss-of-function phenotypes.

The molecular reason for this functional impairment was the deletion of the amphipathic N terminus of Ankrd26, which we demonstrated to insert itself into the hydrophobic part of the membrane, as demonstrated by its insensitivity to suppression of electrostatic interactions and by mutagenesis of hydrophilic and hydrophobic amino acid residues, respectively. Ankrd26 thus shares some properties with ankycorbin, the founding member of the recently suggested superfamily of N-Ank proteins (Wolf et al., 2019). Within the N-Ank superfamily, Ankrd26 phylogenetically belongs to a mostly human or primate-specific subfamily, whose functions are mostly completely unknown.

Our functional and molecular studies demonstrated that the *ANKRD26* mutant identified in AML (Marconi et al., 2017) is a clear loss-of-function mutant in terms of Ankrd26’s critical role in cellular differentiation and in terms of Ankrd26’s N-Ank module-mediated membrane binding. Impairments in Ankrd26-mediated cellular differentiation and in Ankrd26’s membrane anchoring and the resulting lack of its spatial confinement to the plasma membrane thus seem to be major aspects associated with thrombocytopenia and with malignancies, such as AML.

Ankrd26 also was reported to be involved in the organization of distal appendages of parent centrioles (Evans et al., 2021; Burigotto et al., 2021). Distal appendages are required for basal body docking to the plasma membrane and thereby enable ciliogenesis (Tanos et al., 2013). While the exact role of Ankrd26 in centriolar and/or ciliary functions remains somewhat elusive, as neither ciliogenesis nor centriole duplication was disrupted in cells lacking Ankrd26 (Yan et al., 2020; Evans et al., 2021; Burigotto et al., 2021), at least Ankrd26’s role in gating ciliary entry of receptors when additionally the centrosomal protein TALPID3 was knocked down (Yan et al., 2020) may relate to the membrane association of Ankrd26 we identified.

It however needs to be emphasized that Ankrd26’s localization is not restricted to centrosomes. Instead, Ankrd26 was shown to be present throughout the soma of cells (Acs et al., 2015). Also our fractionation data showed a significant portion of endogenous Ankrd26 in fraction S2 representing predominantly soluble proteins. Additionally, our fractionation analyses of endogenous as well as of GFP-tagged Ankrd26, our immunofluorescence studies and our electron microscopical detections of endogenous Ankrd26 clearly demonstrated that Ankrd26 is in fact present at wide areas of the plasma membrane. As tumor cells frequently show dysregulated centriole numbers (either loss or extra copies) (Nigg and Holland, 2018), it seems possible that ciliary length alterations described in mice expressing a β-galactosidase gene trap fusion of Ankrd26 may in fact mostly relate to altered signaling pathways. In line with this, these Ankrd26 mutant mice were obese and much larger than WT mice and for some reasons showed increased Akt, mTOR, insulin receptor and insulin growth factor 1 receptor signaling (Bera et al., 2008).

Our analyses of Ankrd26-related pathomechanisms demonstrated that not only Ankrd26’s membrane interaction is an important aspect but that Ankrd26 also is able to self-associate and that an Ankrd26 mutant lacking the coiled coil domain-containing C terminal part of Ankrd26 is unable to rescue any of the Ankrd26 loss-of-function defects in cellular differentiation of neuroblastoma cells. Our data hereby are fully in line with the detection of Ankrd26 clusters at the plasma membranes of freeze-fractured cells. The small distances of immunogold labels inside of these clusters hereby were in the theoretical extension range of two neighboring Ankrd26 proteins and/or of two Ankrd26-detecting probes. Ankrd26 self-association into multimeric clusters at the plasma membrane suggests that Ankrd26 can provide signaling and/or organizational hubs at cellular membranes.

RET is a plasma membrane-localized receptor tyrosine kinase connected to a variety of down-stream signaling pathways controlling cellular proliferation (Morandi et al., 2011; Shaw et al., 2013; Mulligan, 2014; Salvatore et al., 2021). Plasma membrane anchoring of RET is mediated by the RET transmembrane domain lacking in the Ankrd26-RET fusion. Furthermore, ligand-induced multisubunit receptor complex formation, which is normally mediated by extracellular interactions, is required for efficient activation of RET signaling (Morandi et al., 2011; Shaw et al., 2013; Mulligan, 2014; Salvatore et al., 2021). Downstream signaling seems to in part also require endocytosis and endosomal trafficking of activated RET (Richardson et al., 2006). Since the RET piece of the Ankrd26-RET fusion solely represents the intracellular kinase domain of RET (Staubitz et al., 2019), important aspects in RET functions seem to be disrupted in the RET fragment included in the Ankrd26-RET fusion product found in papillary thyroid carcinoma patients. Our results show that the papillary thyroid carcinoma-associated Ankrd26-RET fusion obviously does have both of these capabilities, membrane anchoring and clustering. Efficient membrane targeting was ensured by Ankrd26’s intact N-Ank module and self-association surprisingly was maintained by the fragment of the coiled coil domain of Ankrd26 still present in the papillary thyroid carcinoma-associated Ankrd26-RET fusion mutant.

The expression of the carcinoma-associated Ankrd26-RET fusion mutant led to an aberrantly strong increase of ERK1/2 activation. This is in line with the ERK cascade functioning in cellular proliferation, differentiation, and survival, and its inappropriate activation being a common occurrence in human cancers. The ERK cascade can be triggered by RET activation in different cancers (Shaw et al., 2013; Liu et al., 2021). In line, we observed that the papillary thyroid carcinoma-associated Ankrd26-RET fusion showed a strong autophosphorylation of the RET kinase domain. RET signaling, which is thought to be organized in lipid rafts as signaling hubs (Tansey et al., 2000), thus seems to be successfully mimicked by Ankrd26-RET and by the Ankrd26 properties we identified. As a membrane binding protein, Ankrd26 mimics the membrane association of RET in a very effective, probably constitutive manner when a fragment of Ankrd26 including the membrane-binding N-Ank module is fused to the RET kinase domain.

We did not only observe that the Ankrd26 part of the Ankrd26-RET fusion can mimic RET functions successfully. Strikingly, we determined a very high autophosphorylation of the papillary thyroid carcinoma-associated Ankrd26-RET fusion mutant that exceeded that of full-length RET. Similarly, the activity of the ERK signaling cascade was much higher for expression of the Ankrd26-RET fusion mutant than for expression of wild-type and full-length RET. Therefore, the Ankrd26-RET fusion mutation represents a completely derailed RET signaling because it was both grossly exaggerated and furthermore completely decoupled from any extracellular ligand binding.

The additional analyses of Ankrd26-RET fusion mutants deficient for selective Ankrd26 properties, which we identified in our studies, highlighted that both effective membrane anchoring as well as confined spatial organization in small subdomains in the plasma membrane add to the grossly exaggerated RET autophosphorylation. Interestingly, the strongly elevated RET kinase domain autophosphorylation caused by fusion with a palmitoylation site-containing GFP demonstrated that the pathomolecular insights obtained by studying the functions of Ankrd26 and its fragment fused to RET can be generalized and also lead to insights into RET signaling and associated pathomechanisms. Constitutive plasma membrane-association and the formation of RET kinase domain-enriched signaling hubs at the plasma membrane seem to be sufficient for grossly exaggerated RET autophosphorylation and its decoupling from extracellular ligand cues.

Taken together, Ankrd26, thus is a member of the novel N-Ank superfamily, that is able to self-associate, to insert itself into the cytosolic membrane leaflet of the plasma membrane by amphipathic interactions and to shape membranes into convex membrane topologies by the use of its ankyrin repeat array. The resulting Ankrd26-mediated organizational platforms seem to be of utmost importance for cellular differentiation processes and signaling pathways originating from the plasma membrane, which, if derailed, lead to cancer-associated pathomechanisms involving the Ankrd26 properties we identified.

## Material and Methods

### DNA constructs

Plasmids encoding human Ankrd26-GFP were cloned by PCR using EST clone 6830677 (NM_014915.2) as template and subcloned into pEGFP-N3 (Clontech) using KpnI und BamHI as restriction sites. Human Ankrd26 deletion mutants were generated by PCR introducing appropriate stop codons and restriction sites. These mutants included the Ankrd26 N-Ank module (Ankrd26^N-Ank^- GFP) (aa1-217), an N terminally shortened version only comprising the ankyrin repeats, Ankrd26^Ank^- GFP (aa45-217), Ankrd26^N-Ank^°^AML^-GFP (aa78-217), Ankrd26°^AML^-GFP (aa78-1710), Ankrd26^490-1710^-GFP, Ankrd26^1-1405^-GFP and Ankrd26^490-1405^-GFP as well as two different sets of Ankrd26 point mutants in the amphipathic N terminus generated by two different mutation-introducing forward primers. For full primer list, please see **Table 1**.

**Table 1.**
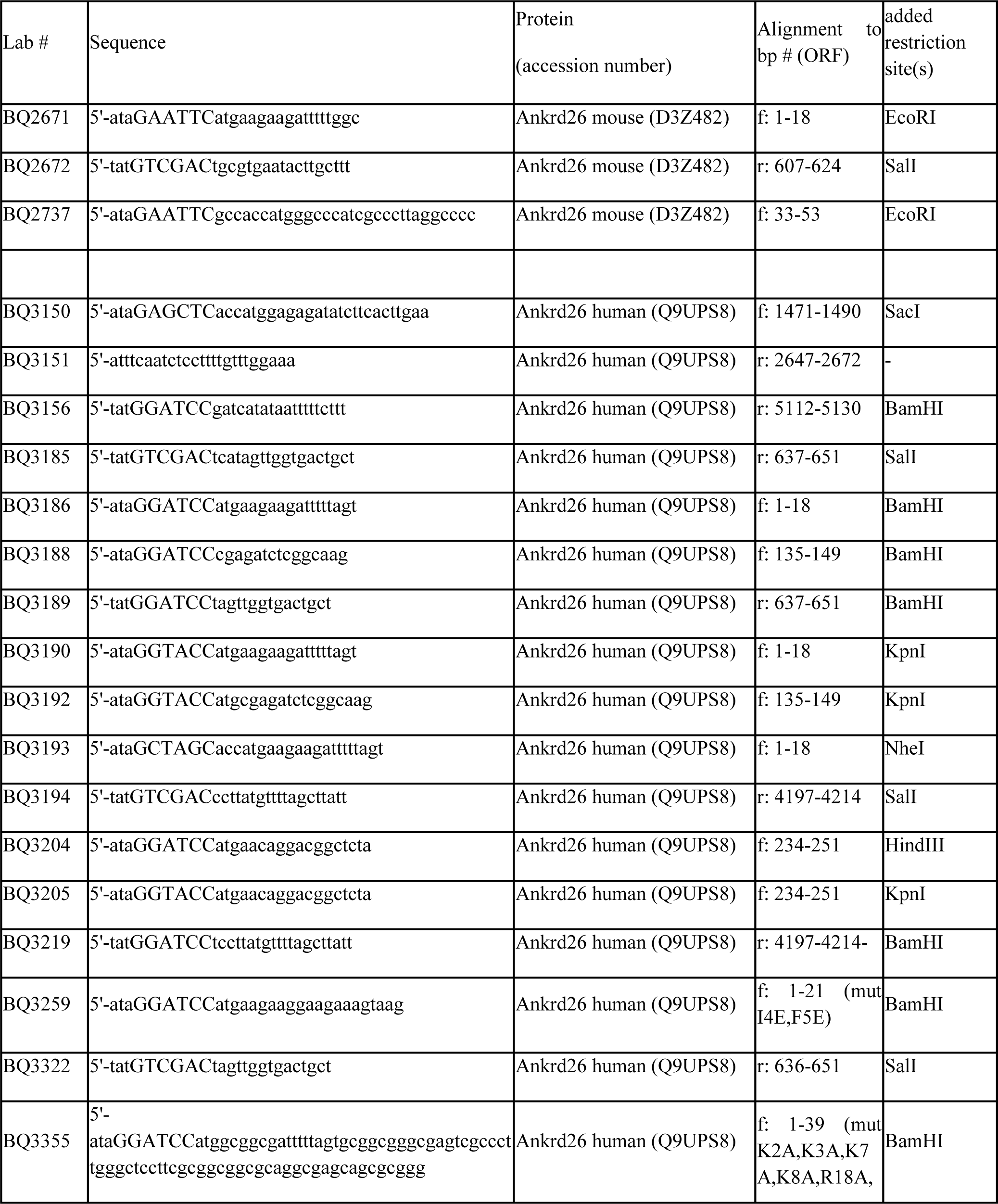
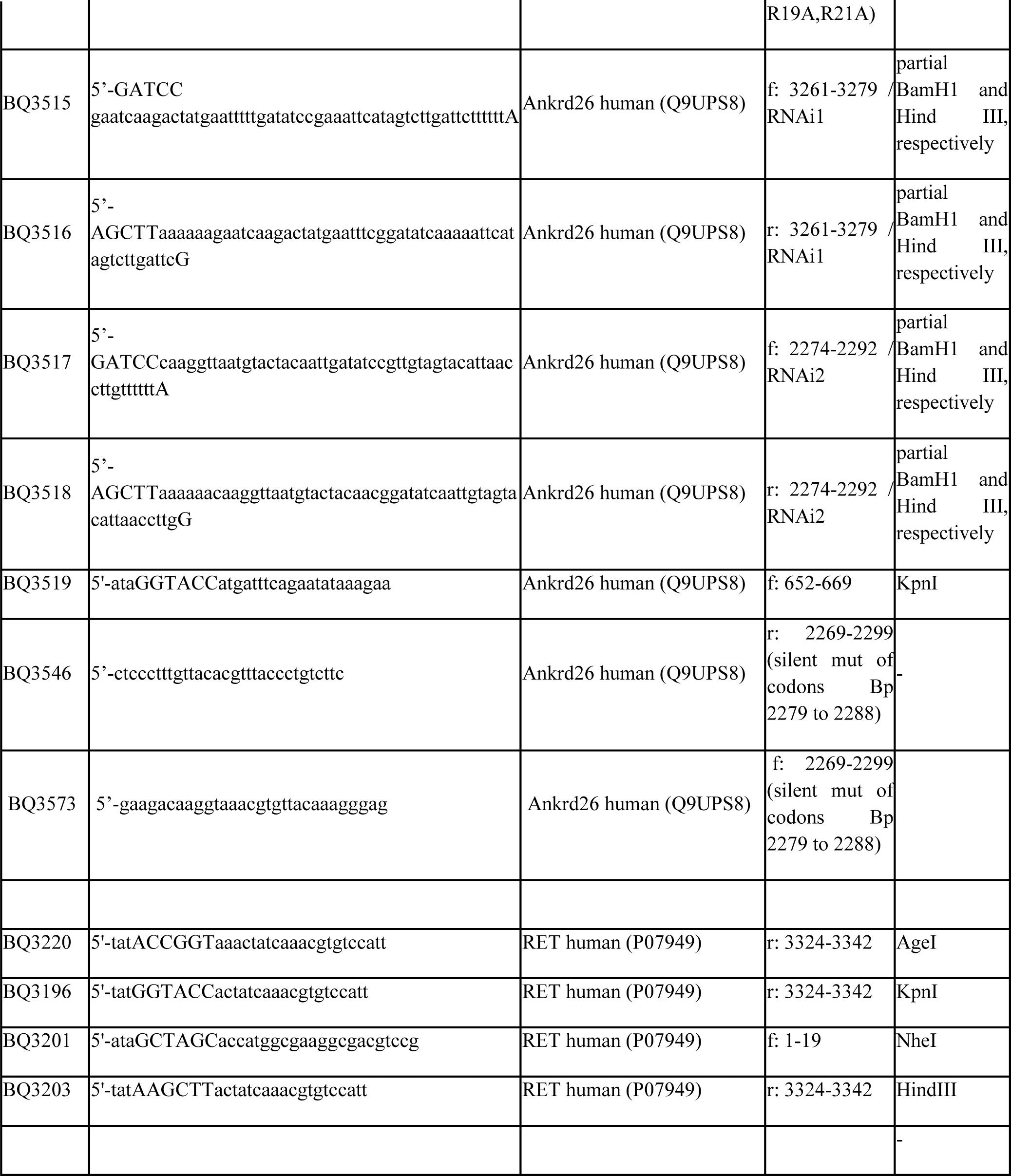
List of primers used. (Partial) restriction sites added are in capital letters. Abbreviations: f, forward; r, reverse. Accession numbers, UniProt.

Mouse Ankrd26^N-Ank^-GFP (aa1-208) and an N terminally deleted construct (Ankrd26^Ank^-GFP (aa11- 208) were cloned from cDNA prepared from brain of 8 weeks old mice. RNA isolation and reverse transcription PCRs were done according to procedures described previously (Haag et al., 2012; Haag et al., 2018). For primers used see **Table 1**.

Plasmids encoding for GST-fused, WT and mutant Ankrd26 N-Ank modules were generated by cloning into pGEX-6P-1 (GE Healthcare).

RNAi tools against Ankrd26 were generated by annealing phosphorylated primers targeting established RNAi sites (Yan et al., 2020) and insertion into pRNAT H1.1-GFP. For primers used please see **Table 1**. A corresponding pRNAT vector expressing a scrambled RNAi sequence served as control (Pinyol et al., 2007).

An RNAi-insensitive Ankrd26-GFP (Ankrd26*-GFP) was generated by introducing several silent mutations into the RNAi2 site (bp 2279 to 2288). The mutations were initially introduced into the Ankrd26-GFP plasmid. For primers used please see **Table 1**. The resulting Ankrd26*-GFP sequence was then used to replace the GFP-encoding sequence in the pRNAT H1.1-based RNAi2 plasmid (NheI/blunted NotI-cut inset into NheI/SmaI-digested RNAi2).

RNAi2 rescue plasmids coexpressing either the AML-associated Ankrd26 mutant (Ankrd26*°^AML^- GFP; i.e. Ankrd26^Δ1-77^-GFP) or Ankrd26*^1-217^-GFP instead of full-length Ankrd26* were generated by subcloning from the respective GFP-N vectors into the RNAi2 plasmid using a similar strategy.

Human RET-GFP and RET^712-1114^-GFP were cloned by PCR using a cDNA clone (GenScript Biotech; NM_020975.6) as template. The PCR products were inserted into pEGFP-N1. For full primer list, please see **Table 1**.

Human Ankrd26^1-1405^ and RET^712-1114^ were fused by complementing Ankrd26^1-1405^ with RET^712-1114^ using the internal AgeI site of the RET-encoding DNA sequence to yield a fusion protein with an amino acid sequence identical to the Ankrd26-RET fusion identified in papillary thyroid carcinoma patients (Staubitz et al., 2019). Ankrd26-RET fusions with mutated Ankrd26 portions were generated by subcloning (Ankrd26^490-1405^-RET^712-1114^; Ankrd26^45-1405^-RET^712-1114^) and by cloning and fusing the Ankrd26 N-Ank sequence to RET^712-1114^ (Ankrd26^1-217^-RET^712-1114^), respectively. For primers, see **Table 1**.

Correct cloning by PCR was verified by sequencing in all cases.

The vector expressing PM-targeted, farnesylated monomeric Cherry (CherryF) was originally provided by M. Korte (TU Braunschweig, Germany) and has been described before to specifically outline the plasma membrane (Hou et al., 2015; Izadi et al., 2018; Wolf et al., 2019).

The vector expressing palmitoylated GFP used to generate PM-GFP-RET^712-114^ by subcloning has been described previously (Dharmalingam et al., 2009).

### Antibodies

Rabbit anti-Ankrd26 antibodies (RRID:AB_867671 and ab183846) and anti-Cherry (RRID:AB_2571870) antibodies were from Abcam and monoclonal mouse anti-GFP antibodies were from Covance (JL-8; RRID AB_10013427). Monoclonal mouse anti-β-actin antibodies were from Sigma (RRID:AB_476744). Mouse monoclonal anti-RET antibodies were from Santa Cruz (6E4C4; RRID:AB_2269604). Rabbit anti-pRET antibodies were from Cell Signaling Technology (pY905; RRID:AB_2179887) and from Abcam (pY1015; RRID:AB_1524311), respectively. Rabbit anti- p44/42 MAPK (ERK1/2) antibodies (RRID:AB_330744) and monoclonal mouse anti-phospho-p44/42 MAPK antibodies (pERK1/2; pT202/pY204) (RRID:AB_331768) were from Cell Signaling.

Secondary antibodies used included DyLight800-conjugated goat anti-rabbit and anti-mouse antibodies (RRID:AB_141725 and RRID:AB_1965956) from Thermo Fisher Scientific. Further secondary antibodies coupled to AlexaFluor680, were from LI-COR Bioscience (anti-rabbit, RRID:AB_2535758; anti-mouse, RRID:AB_1965956). EM grade 10 nm gold anti-rabbit conjugates were from BBI Solutions (EM.GAR10/2).

### Purification of recombinant proteins and tag cleavage

GST-tagged fusion protein purification and PreScission Protease (GE Healthcare) cleavage of pGEX- 6P-1-encoded GST-fusion proteins was performed at 4°C during dialysis against HN buffer (150 mM NaCl, 2 mM DTT, 20 mM HEPES pH 7.4) overnight essentially as described (Seemann et al., 2017; Wolf et al., 2019).

Cleaved off GST and putatively remaining uncut GST-fusion proteins were subsequently removed by affinity purification with glutathione-agarose (Antibodies-Online) and proteins of interest were collected as flow-through by centrifugation.

Protein concentrations were determined by the Bradford method. Successful proteolytic cleavage and protein integrity were verified by SDS-PAGE and Coomassie staining.

### Liposome preparation and sizing

Large unilamellar vesicles (LUVs; liposomes) were prepared from Folch fraction I lipids (Sigma Aldrich) essentially as described (Koch et al., 2011). The mean diameters of LUV preparations can differ in independent liposome preparations. Therefore, all size determinations and liposome size- changing assays were always performed with full set of conditions and controls in each assay.

Small unilamellar vesicles (SUVs) were generated by sonication of liposomes, as described (Zobel et al., 2015). Briefly, liposomes were sonicated 3 times (50 s each, 1 min pause in between) on ice using an ultrasonicator UP50H (Dr. Hielscher). SUVs prepared under these conditions have a mean diameter of about 50 nm, as determined by quantitative electron microscopical examinations (Wolf et al., 2019).

### Liposome cosedimentations (membrane binding, salt resistance, curvature sensing of proteins)

Liposome coprecipitation assays were essentially done as described previously (Wolf et al., 2019). In brief, 4 µM of untagged WT Ankrd26 N-Ank modules or to be tested mutant proteins, respectively, in 25 µl HN buffer were prespun for 5 min at 200000xg to remove putative precipitates. The supernatant was transferred to a fresh tube and incubated with 50 µg of Folch fraction I liposomes (2 mg/ml) for 30 min at RT. Liposomes were pelleted by 200000xg centrifugation (20 min, 28°C). The resulting supernatants (S) were collected immediately. The pellets (P) were resuspended in volumes equal to the supernatant. All fractions were then analyzed by SDS-PAGE and Coomassie staining. Gels were visualized with a LI-COR Odyssey imager and the lane intensities quantified using the LI-COR Odyssey software (LI-COR Bioscience).

The amount of protein coprecipitated with liposomes was expressed as percent of total protein after subtraction of any putative unspecific background of precipitation in HN buffer control, i.e. liposome binding [%] = [((P/(S+P)) Sample - (P/(S+P)) HN buffer control)] x 100.

Salt extraction experiments were essentially performed as described above, except that 5 min prior to centrifugation at 200000xg the final NaCl concentration was increased from 150 mM (HN buffer) to 200 mM and 250 mM, respectively. Liposome binding in relation to the mean of the respective binding in 150 mM NaCl set to 100%.

Curvature sensing activities were analyzed by slightly modified cosedimentation assays using LUVs versus SUVs, as essentially described previously (Wolf et al., 2019). Due to decreased sedimentation efficiencies of SUVs and the accordingly lose pellets, only the upper 35 µl of the 200000xg centrifugation supernatant were collected and supplemented with 4x SDS-PAGE sample buffer yielding 47 µl of sample. For direct comparison, the pellet-containing fraction was also resuspended in a final volume of 47 µl. Both samples were then analyzed by SDS-PAGE and Coomassie staining. The signals were quantified and, as in LUV binding experiments described above, traces of protein precipitating in control incubations without any liposomes added were subtracted from the data of liposome binding as unspecific background. The data were then expressed as deviation from mean LUV binding.

### Culturing, transfection, immunostaining and fluorescence microscopy of cells

Culturing of HEK293 (RRID:CVCL_0045) and HeLa cells (RRID:CVCL_0030) and their transfection using TurboFect (Thermo Fisher Scientific) was essentially done as described (Kessels et al., 2001; Haag et al., 2012).

SK-N-SH cells (RRID:CVCL_0531) were cultured on poly-D-lysine-coated coverslips in 24-well plates and were differentiated by adding 10 µM all-trans retinoic acid (Sigma; R2625) on day 2 and conducting a media exchange to brain-derived neurotrophic factor (BDNF)-containing media (DMEM with 50 ng/ml BDNF, PreproTech; 080861) at day 5. At day 8, cells were transfected using TurboFect as described before (Haag et al., 2018; Izadi et al., 2021). Prior to fixation two days later, cells were washed with PBS and incubated with 0.05% (w/v) tetramethylrhodamine-coupled wheat germ agglutinin (WGA) (Molecular Probes) in PBS for 30 min at RT. After washing with PBS, cells were fixed and stained with Alex Flour 647-coupled phalloidin (Molecular Probes) and DAPI essentially as described before (Izadi et al., 2018).

Images were recorded as z-stacks using a Zeiss AxioObserver.Z1 microscope equipped with an ApoTome for pseudo-confocal image recording (Zeiss), Plan-Apochromat 63x/1.4, and 40x/1.3 objectives (Zeiss) and an AxioCam MRm CCD camera (Zeiss).

Digital images from Zeiss microscopes were recorded by ZEN2012 (PRID:SCR_013672). Image processing was done by Adobe Photoshop (RRID:SCR_014199).

### Quantitative morphometric analyses of differentiated SK-N-SH neuroblastoma

For quantitative analyses, 2-3 independent coverslips per condition per assay and transfected cells of 2-4 independent preparations were analyzed based on the WGA and F-actin staining of the cells. Transfected cells were sampled systematically on each coverslip.

Phenotypical analyses were conducted using IMARIS 8.4 software (RRID:SCR_007370) to construct a 3D-morphological trace (“filament”) of each cell. Initial phenotype screening analyses were based on tracking protrusion number and protrusion length from the cell perimeter using the following IMARIS 8.4 software settings: thinnest diameter, 0.2 µm; minimum protrusion size, 10 µm; disconnected points, 2 µm.

Subsequent analyses were done by a different experimenter to validate phenotypes independently. These analyses addressed both gain- and loss-of-function phenotypes and employed IMARIS 8.4 software settings designed to also cover smaller cellular protrusions. Using the IMARIS start seed point as start of protrusions, the IMARIS 8.4 software settings were as follows: thinnest diameter, 0.2 µm; minimum protrusion size, 5 µm; disconnected points, 2 µm.

Immunopositive areas that were erroneously spliced by IMARIS or protrusions belonging to different cells as well as points that the software erroneously placed inside of a cell body were manually corrected. Parameters determined were exported and saved as Excel files. Statistical significance calculations were done using GraphPad Prism8 (RRID:SCR_002798) software.

### Freeze-fracturing and TEM of freeze-fracture replica

For determinations of size changes of liposomes by Ankrd26 proteins, 100 µg liposomes were incubated with a final protein concentration of 6 µM Ankrd26 N-Ank module and mutants thereof, respectively, and compared to the results obtained by incubation without protein and with 6 µM GST as unrelated protein control, respectively, in a final volume of 100 µl for 15 min at 37°C. To avoid liposomal aggregation during sample preparation, the suspension was subsequently incubated with 30 µg proteinase K for 25 min at 45°C. Liposomes were centrifuged for 15 min at 200000xg and the pellet was resuspended in 20 µl of the supernatant to obtain a concentrated liposome preparation for freeze-fracture (Beetz et al., 2013).

Freeze-fracturing of liposomes was done using two 0.6 mm high copper profiles with 2 µl of the liposome solution placed between them essentially as described previously (Seemann et al., 2017; Wolf et al., 2019).

Freeze-fracturing of SK-N-SH cells was done as follows: cells were seeded onto collagen-coated (Sigma) sapphire discs in 24-well cell culture plates and differentiated using retinoic acid and BDNF as described above. After a media change at day 8, cells were subjected to freeze-fracturing on day 9. After washing with PBS, cells were covered with 20% (w/v) BSA (in PBS) and a copper head sandwich profile and then immediately plunge-frozen in liquid propane:ethane (1:1) cooled in liquid nitrogen. Sandwiches were placed in a double-replica specimen table cooled by liquid nitrogen that was then transferred into the freeze-fracture machine BAF400T (Leica) cooled to −140°C. Freeze-fracturing was done at ≤10^-6^ mbar.

Immediately after freeze-fracturing, 2 nm platinum/carbon was evaporated onto the samples (angle, 35°). The replica were then stabilized by evaporating a carbon coat of 15-20 nm onto the samples (angle, 90°). The samples were then carefully extracted from the freeze-fracture machine, thawed and floated onto ddH_2_O. Replica were then incubated for 15 min in 3% (v/v) sodium hypochlorite warmed to 50°C, washed twice with ddH_2_O for 10 min, transferred onto uncoated copper grids (300 mesh) and dried.

For immunogold labeling of freeze-fractured replica, the replica were washed three times with PBS and incubated in blocking buffer (1% (w/v) BSA, 0.5% (w/v) fish gelatin, 0.005% (v/v) Tween 20 in PBS) for 30 min at RT, followed by incubation with the primary rabbit antibody (anti-Ankrd26 ab183846 (Abcam); 1:50 in blocking buffer) at 4°C overnight. The next day, unbound antibody was removed by washing the samples with PBS three times and anti-rabbit 10 nm gold-conjugated secondary antibody (1:50) was added for 1.5 h at RT. Afterwards, the samples were washed two times with PBS and fixed with 0.5% (v/v) glutaraldehyde in PBS and washed two times with ddH_2_O for 10 min and transferred onto uncoated copper grids.

Imaging of freeze-fracture replica was done by TEM (EM902A, Zeiss). The electron microscope was operated at 80 keV.

Imaging was done by systematic explorations of the grids. Images were recorded with a CCD camera (TVIPS; EM-Menu 4) and processed with Adobe Photoshop.

### Quantitative EM analyses of liposomes

Diameters of freeze-fractured liposomes were measured using ImageJ (RRID:SCR_003070), as described previously (Wolf et al., 2019). Irregular structures were excluded from analysis.

All TEM studies of liposomes were conducted in a fully blinded manner.

### Quantitative analyses of Ankrd26 immunogold labeling at freeze-fracture replica of SK-N-SH cells

Images were collected by systematic grid explorations. All analyses were done with two independent cell preparations. For determinations of labeling densities, membrane areas for P face, E face and ice were distinguished by their visible topologies and measured using ImageJ.

Cluster analyses were conducted using circular ROIs of 100 nm diameter. In the rare cases of superclusters exceeding the size of 100 nm in diameter, these extended nanodomains were split into 100 nm subclusters and considered as several merged clusters. Gold particles per ROI were counted and single, paired, and clustered particles were analyzed (categories of clustered particles, 3-5 and >5).

### Preparation of membrane-enriched fractions from HEK293 cells

HEK293 cells were either left untreated (fractionations addressing endogenous Ankrd26) or were transfected with GFP, GFP-tagged Ankrd26 and mutants thereof as well as with RET-GFP and mutants thereof in combination with mCherryF as plasma membrane marker. 24 h post transfection, the cells were washed with PBS, harvested, collected by centrifugation (1000xg 5 min, 4°C), resuspended in homogenization buffer (0.32 M sucrose, 5 mM HEPES pH 7.4, 1 mM EDTA) containing protease inhibitor Complete (Roche) and then homogenized by multiple passing through a 0.24 mm syringe.

The resulting homogenates were fractionated essentially as described (Wolf et al., 2019). In brief, the homogenates were subjected to centrifugation at 4°C for 10 min at 1000xg yielding fractions S1 and P1. S1 was then centrifuged further for 20 min at 118000xg. After removal of the supernatant S2, the resulting membrane-containing pellet P2 was resuspended in homogenization buffer und again centrifuged at 11800xg for 20 min resulting in fractions S2’ and P2’. P2’ thus represents a washed membrane-containing fraction P2.

All pellet fractions were resuspended in volumes equal to those of the corresponding supernatants. SDS-PAGE was performed with equal amounts of supernatant and pellet fractions and analyzed by (quantitative) immunoblotting using a LI-COR Odyssey System for fluorescence detection (LI-COR Biosciences GmbH).

### Crosslink studies

Crosslink studies with lysates of HEK293 cells overexpressing different GFP fusion proteins were essentially done as described for endogenous proteins (Kessels and Qualmann, 2006). The cell lysates were incubated in the presence of increasing amounts of the heterobifunctional crosslinker 1-ethyl-3- [dimethylaminopropyl]carbodiimide (EDC) (Pierce) in 20 mM HEPES, pH 7.4, 0.2 mM MgCl_2_, and 2 mM EGTA for 20 min at 30°C. Subsequently, the crosslinking reaction was stopped by adding 4x SDS-PAGE sample buffer and incubating the samples at 95°C for 5 min. The samples then were analyzed by immunoblotting (tank blotting to PVDF membranes).

### ERK and RET activity determinations by quantitative immunoblotting of pERK1/2 and ERK1/2 as well as by quantitative determinations of RET Y905 and RET Y1015 phosphorylations

HEK293 cells were transfected with plasmids encoding for different GFP-tagged proteins and harvested 48 h later. Cell lysis was conducted in radioimmunoprecipitation (RIPA) buffer (150 mM NaCl, 10 mM Na_3_PO_4_ pH 7.2, 1% (w/w) Nonidet P-40, 0.5% (w/v) sodium desoxycholate, 2 mM EDTA, 50 mM NaF, protease inhibitor cocktail Complete EDTA-free (Roche), phosphatase inhibitor cocktail PhosSTOP (Roche)) for 20 min at 4°C and by sonication (5 pulses of each 1 s). The lysates were centrifuged at 10000xg for 10 min at 4°C. The supernatants were incubated at 95°C with SDS-PAGE sample puffer for 5 min and were analyzed by immunoblotting. Anti-GFP and anti-RET antibodies, respectively, were used to determine the expression levels of the proteins to be analyzed. Anti-pERK1/2 antibodies (anti-T202/Y204) and anti-pRET antibodies (anti-pY905 and anti-pY1015, respectively) served to determine the phosphorylation status of ERK1/2 and the autophosphorylation of RET, respectively. The immunosignals were quantitatively analyzed using fluorescent antibodies, a LI-COR Odyssey System and Image Studio Lite V5.2 software (RRID:SCR_015795) for band intensity determinations.

### In silico analyses

Amphipathic helical wheel representations were done using HELIQUEST (Gautier et al., 2008). The ankyrin repeat array of human Ankrd26 was modeled in two rotational views employing https://swissmodel.expasy.org/viewer/ngl/. Coiled coil predictions were done with COILS (RRID:SCR_008440; https://www.expasy.org/resources/coils). Alignments were done using Clustal Omega (RRID:SCR_001591; https://www.ebi.ac.uk/Tools/msa/clustalo/).

### Data reporting and statistical analyses

No statistical methods were used to predetermine sample size. All TEM experiments with liposomes were randomized and the investigators were blinded to allocation during experiments and outcome assessment.

No outlier suggestions were computed. All quantitative evaluation data points were taken into account and averaged to fully represent biological and technical variabilities. Wherever useful, the bar plots in the figures have been overlayed with dot plots showing all individual data points measured.

Except distribution analyses (**Figure 4K; Figure 9N**), all quantitative data shown represent mean and SEM. Statistical analyses were done using GraphPad Prism software. Statistical significances were marked by \**P*<0.05; \*\**P*<0.01; \*\*\**P*<0.001; \*\*\*\**P*<0.0001 throughout.

Determinations of liposome binding, SUV versus LUV preference in binding as well as differences in liposome diameters were analyzed for significances by either Mann-Whitney-U-test or two-tailed, unpaired Student’s t-test (depending on normal data distribution; comparisons of two conditions) or by two-way ANOVA or by Kruskal Wallis (multiple conditions), respectively.

Quantitative data from extractions of liposome-bound proteins with salt were tested using one-way ANOVA with Tukey post hoc test.

Comparisons of anti-Ankrd26 labeling densities were also done using one-way ANOVA with Tukey post hoc test.

Phenotypes of Ankrd26 deficiency compared to scrambled RNAi were analyzed for statistical significances by two-tailed, unpaired Student’s t-tests. The results of RNAi rescue experiments with

Ankrd26 mutants instead of full-length Ankrd26 were analyzed by one-way ANOVA with Tukey post hoc test. Similarly, overexpression experiments with Ankrd26-GFP and the AML-associated mutant thereof in comparison to GFP were analyzed for statistical significances using one-way ANOVA with Tukey post hoc test.

Determinations of pY905 and pY1015 autophosphorylation levels of the RET kinase domain included in different fusion proteins and mutants were tested by Bartlett’s test and Tukey post hoc test. Examinations of ERK1/2 activation levels by quantitative Western blot analyses were tested using one-way ANOVA with Tukey post hoc test.

### Data availability

This study includes no data deposited in external repositories. All data generated or analyzed during this study are included in this published article (and its supplementary information files). Numerical data of all quantitative analyses are provided as Supplementary Data.

## Acknowledgments

We thank D. Wolf for valuable initial observations. We furthermore thank B. Schade, M. Roeder, S. Berr, K. Gluth and S. Linde for technical assistance, E. Seemann for EM instructions, advice and practical help and M. Westermann for access to freeze-fracturing and TEM.

This work was supported by *DFG* grants to M.M.K. (KE685/7-1) and B.Q. (QU116/9-1).

## Supplementary Information

**Figure 1-figure supplement 1.**
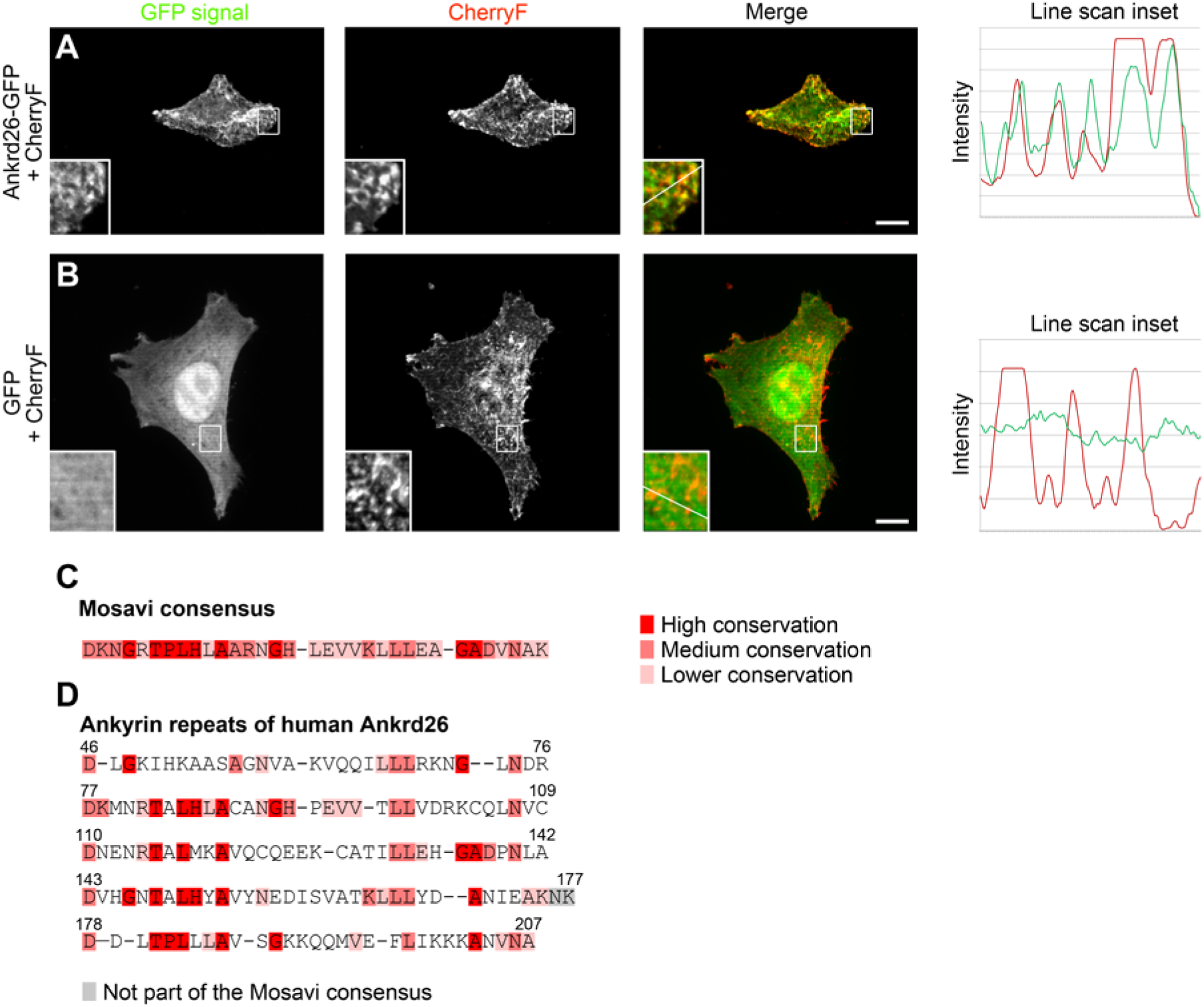
Human Ankrd26 is a membrane-binding, five ankyrin repeat- containing protein. **(A,B)** Maximum intensity projections (MIPs) of Ankrd26-GFP and GFP in HeLa cells cotransfected with the plasma membrane marker CherryF shown in **Figure 1C,D** with intensity plots of the red and green fluorescent channel along randomly positioned lines marked in the insets, which represent magnifications of boxed areas. Note that in **A** both fluorophores show synchronous intensity changes (coenrichment), whereas the control shown in **B** does not show any correlation. Bars, 10 µm. **(C)** Ankyrin repeat consensus according to Mosavi et al. (2004). **(D)** The five suggested ankyrin repeats of human Ankrd26 (according to UniProt) and their overlap with the Mosavi consensus. Residues of high, medium and lower conservation in the Mosavi consensus are highlighted in different shades of red, as indicated. Grey marks residues that are not following the consensus and are located between he ankyrin repeats of Ankrd26 (included only to report a continuous sequence of the complete ankyrin repeat array).

**Figure 1-figure supplement 2.**
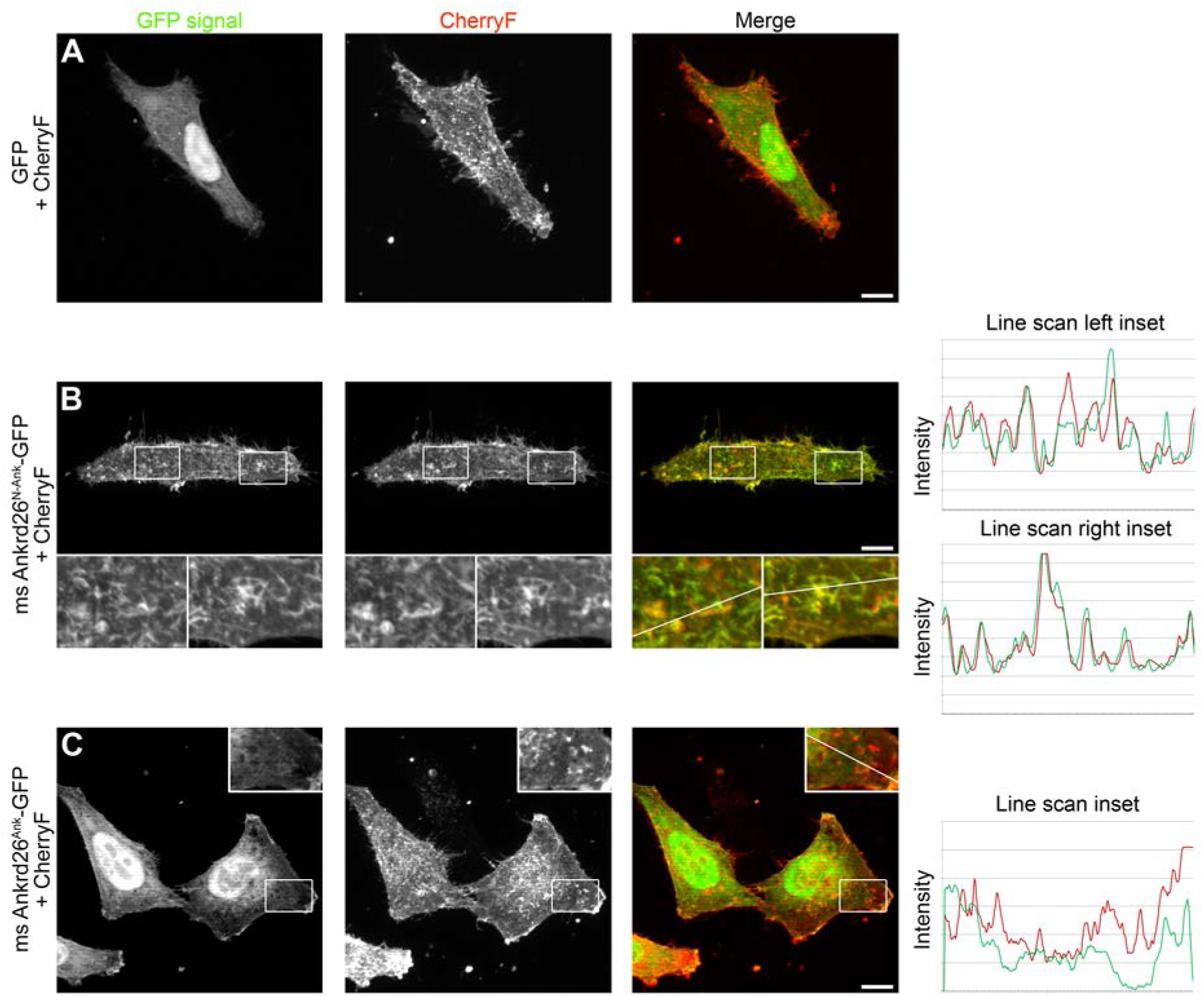
Mouse Ankrd26^N-Ank^ but not a mutant lacking the Ankrd26 N terminus colocalizes with the plasma membrane marker CherryF. **(A-C)** MIPs of GFP **(A)**, mouse Ankrd26^N-Ank^-GFP **(B)** and mouse Ankrd26^Ank^-GFP **(C)**, respectively, in HeLa cells cotransfected with CherryF to mark the plasma membrane. Note the spatial overlap of ms Ankrd26^N-Ank^ with CherryF. Insets in **B** and **C** represent enlargements of boxed areas. Lines in insets mark positions of intensity analyses (see line plots of the red and green fluorescent channel). Bars, 10 µm.

**Figure 3-figure supplement 1.**
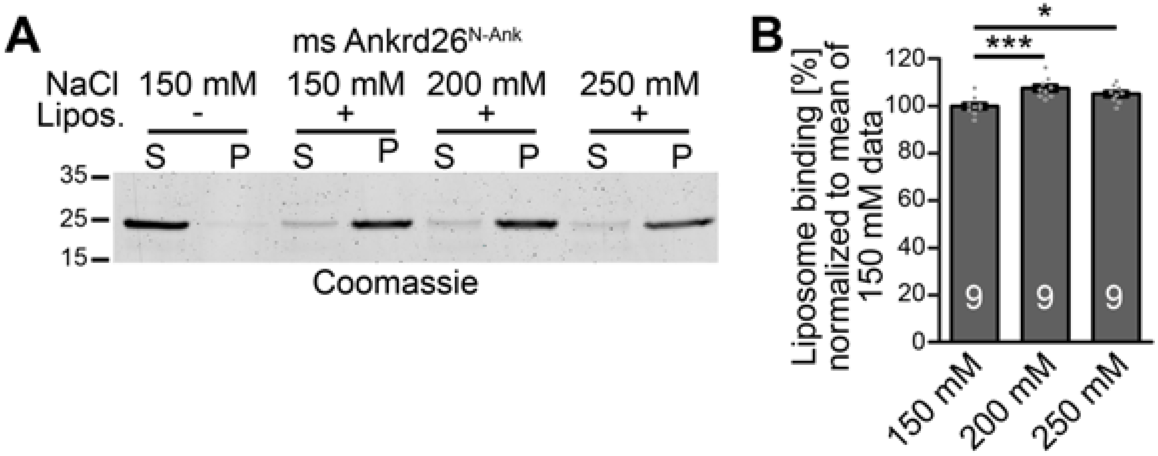
Mouse Ankrd26 binds to membranes via non-electrostatic interactions critically relying on its amphipathic N terminal structure. **(A,B)** Attempts of extraction of liposome-bound mouse Ankrd26^N-Ank^ using increasing salt concentrations. **(A)** Representative image of a Coomassie-stained SDS-PAGE gel. **(B)** Quantitative analyses (**B**) confirming that also the liposome binding of ms Ankrd26^N-Ank^ is fully salt-resistant. Data represent mean±SEM. Bar plots with individual data points (dot plots). n=9 experiments each; one- way ANOVA with Tukey post hoc test. \**P*<0.05; \*\*\**P*<0.001.

**Figure 4-figure supplement 1.**
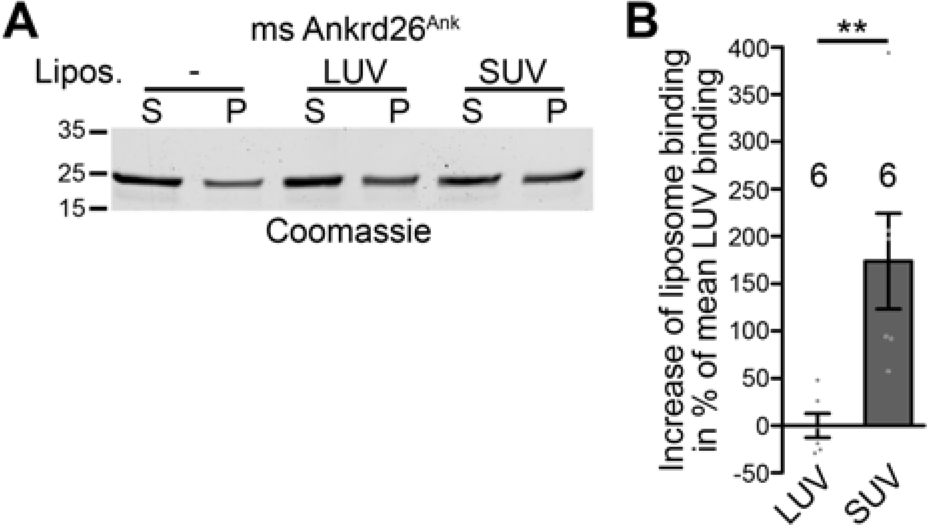
Mouse Ankrd26 recognizes membrane curvatures by its ankyrin repeat array. **(A,B)** The ankyrin repeat array of mouse Ankrd26 prefers the highly curved membranes of SUVs over the more shallowly curved membrane surfaces of LUVs, as demonstrated by comparative coprecipitations **(A)** and quantitative analyses thereof **(B)**. Data represent mean±SEM. Bar plots with individual data points (dot plots). n=6 experiments each; two tailed, unpaired Student’s t-test. \*\**P*<0.01.

**Figure 8-figure supplement 1.**
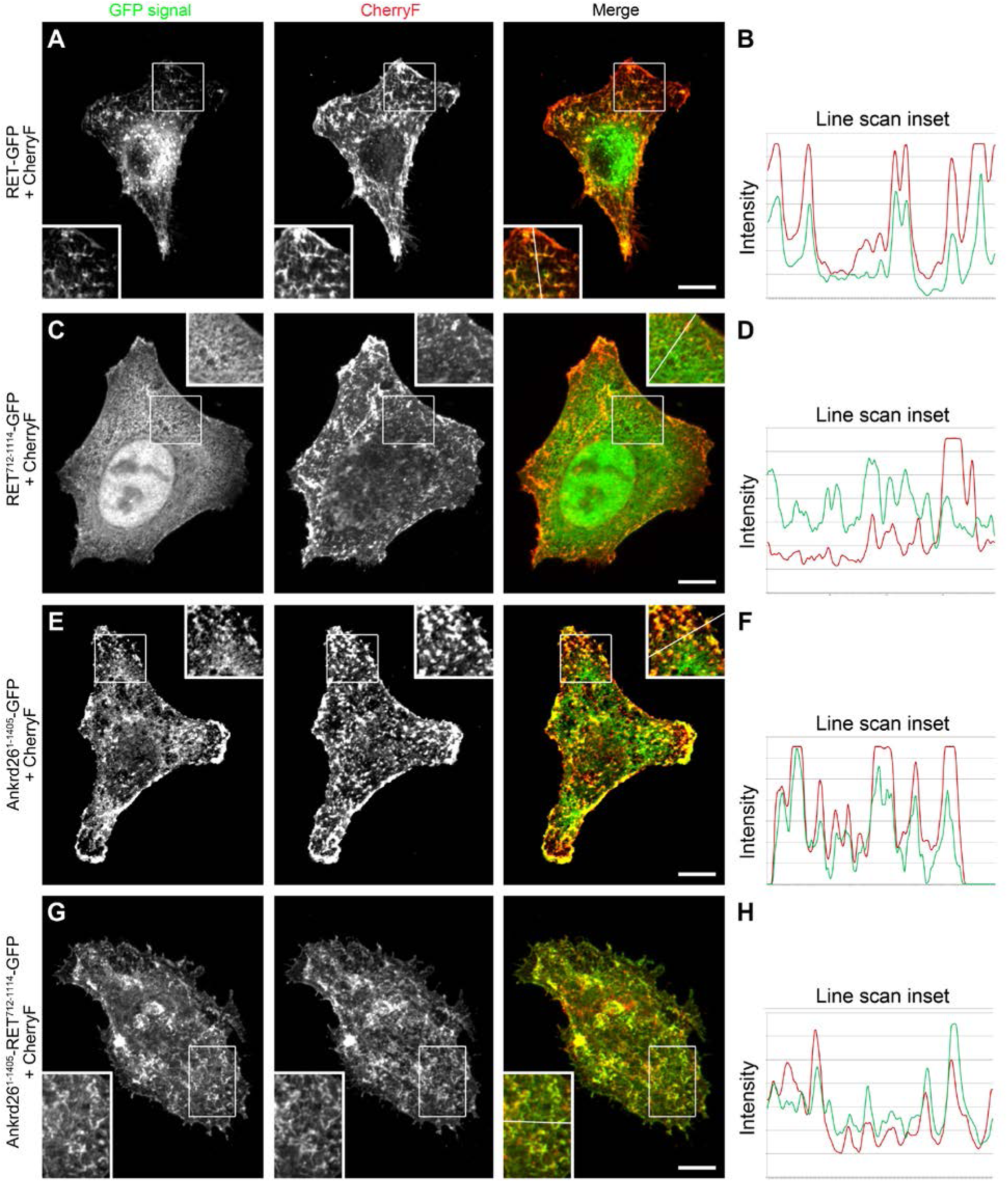
The papillary thyroid carcinoma-associated fusion of Ankrd26 and RET exhibits a membrane localization mediated by the included Ankrd26 portion. **(A-H)** MIPs of indicated proteins in CherryF-cotransfected HeLa cells **(A,C,E,G)**, as in **Figure 8**, supplemented with fluorescence intensity plots (arbitrary units) for the red and the green fluorescence channel **(B,D,F,H)** along the lines indicated in the insets (see line added), which are magnifications of boxed areas. Bars, 10 µm.

**Figure 11-figure supplement 1.**
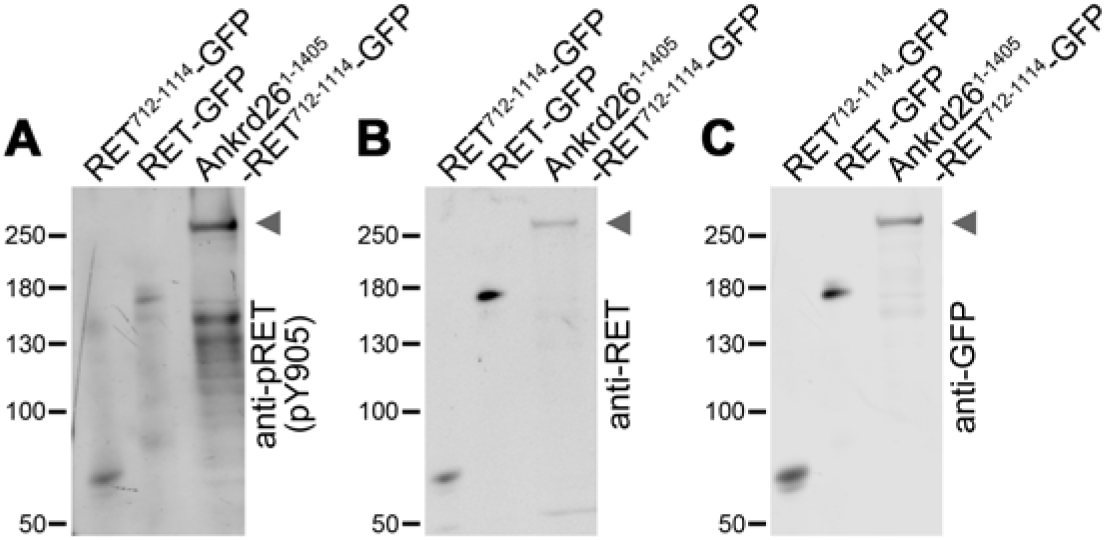
The papillary thyroid carcinoma-associated Ankrd26-RET fusion mutant exhibits a strongly increased RET Y905 autophosphorylation. **(A-C)** Representative images of lysates of HEK293 cells transfected with GFP-tagged RET^712-1114^, full-length RET (RET^1-1114^) and the papillary thyroid carcinoma-associated fusion Ankrd26^1-1405^- RET^712-1114^, respectively, that were immunoblotted with antibodies specifically detecting pRET (pY905) **(A)**, with antibodies against the C terminal domain of RET **(B)** and with anti-GFP antibodies **(C)**, respectively. Arrowheads mark the positions of the highly Y905 phosphorylated Ankrd26^1-1405^- RET^712-1114^-GFP.

## References

Acs P, Bauer PO, Mayer B, Bera T, MacAllister R, Mezey E, Pastan I. A novel form of ciliopathy underlies hyperphagia and obesity in Ankrd26 knockout mice. Brain Struct Funct (2015) 220: 1511–1528.

Arni S, Keilbaugh SA, Ostermeyer AG, Brown DA. Association of GAP-43 with detergent-resistant membranes requires two palmitoylated cysteine residues. J Biol Chem (1998) 273: 28478–28485.

Beetz C, Koch N, Khundadze M, Zimmer G, Nietzsche S, Hertel N, Huebner AK, Mumtaz R, Schweizer M, Dirren E, Karle KN, Irintchev A, Alvarez V, Redies C, Westermann M, Kurth I, Deufel T, Kessels MM, Qualmann B, Hübner CA. A spastic paraplegia mouse model reveals REEP1- dependent ER shaping. J Clin Invest (2013) 123: 4273–4282.

Bera TK, Zimonjic DB, Popescu NC, Sathyanarayana BK, Kumar V, Lee B, Pastan I. POTE, a highly homologous gene family located on numerous chromosomes and expressed in prostate, ovary, testis, placenta, and prostate cancer. Proc Natl Acad Sci USA (2002) 99: 16975–16980.

Bera TK, Saint Fleur A, Lee Y, Kydd A, Hahn Y, Popescu NC, Zimonjic DB, Lee B, Pastan I POTE paralogs are induced and differentially expressed in many cancers. Cancer Res (2006) 66: 52–56.

Bera TK, Liu XF, Yamada M, Gavrilova O, Mezey E, Tessarollo L, Anver M, Hahn Y, Lee B, Pastan I. A model for obesity and gigantism due to disruption of the Ankrd26 gene. Proc Natl Acad Sci USA (2008) 105: 270–275.

Bluteau D, Balduini A, Balayn N, Currao M, Nurden P, Deswarte C, Leverger G, Noris P, Perrotta S, Solary E, Vainchenker W, Debili N, Favier R, Raslova H. Thrombocytopenia-associated mutations in the ANKRD26 regulatory region induce MAPK hyperactivation. J Clin Invest (2014) 124: 580–591.

Burigotto M, Mattivi A, Migliorati D, Magnani G, Valentini C, Roccuzzo M, Offterdinger M, Pizzato M, Schmidt A, Villunger A, Maffini S, Fava LL. Centriolar distal appendages activate the centrosome-PIDDosome-p53 signalling axis via ANKRD26. EMBO J (2021) 40: e104844.

Cerami E, Gao J, Dogrusoz U, Gross BE, Sumer SO, Aksoy BA, Jacobsen A, Byrne CJ, Heuer ML, Larsson E, Antipin Y, Reva B, Goldberg AP, Sander C, Schultz N. The cBio cancer genomics portal: an open platform for exploring multidimensional cancer genomics data. Cancer Discov (2012) 2: 401–404.

Dharmalingam E, Haeckel A, Pinyol R, Schwintzer L, Koch D, Kessels MM, Qualmann B. F-BAR proteins of the syndapin family shape the plasma membrane and are crucial for neuromorphogenesis. J Neurosci (2009) 29: 13315–13327.

Encinas M, Iglesias M, Liu Y, Wang H, Muhaisen A, Ceña V, Gallego C, Comella JX. Sequential treatment of SH-SY5Y cells with retinoic acid and brain-derived neurotrophic factor gives rise to fully differentiated, neurotrophic factor-dependent, human neuron-like cells. J Neurochem (2000) 75: 991–1003.

Evans LT, Anglen T, Scott P, Lukasik K, Loncarek J, Holland AJ. ANKRD26 recruits PIDD1 to centriolar distal appendages to activate the PIDDosome following centrosome amplification. EMBO J (2021) 40: e105106.

Fava LL, Schuler F, Sladky V, Haschka MD, Soratroi C, Eiterer L, Demetz E, Weiss G, Geley S, Nigg EA, Villunger A. The PIDDosome activates p53 in response to supernumerary centrosomes. Genes Dev (2017) 31: 34–45.

Gautier R, Douguet D, Antonny B, Drin G. HELIQUEST: a web server to screen sequences with specific alpha-helical properties. Bioinformatics (2008) 24: 2101–2102.

Haag N, Schwintzer L, Ahuja R, Koch N, Grimm J, Heuer H, Kessels MM, Qualmann B. The actin nucleator Cobl is crucial for Purkinje cell development and works in close conjunction with the F- actin binding protein Abp1. J Neurosci (2012) 32: 17842–17856.

Haag N, Schüler S, Nietzsche S, Hübner CA, Strenzke N, Qualmann B, Kessels MM. (2018) The Actin Nucleator Cobl Is Critical for Centriolar Positioning, Postnatal Planar Cell Polarity Refinement, and Function of the Cochlea. Cell Rep (2018) 24: 2418–2431.

Hou W, Izadi M, Nemitz S, Haag N, Kessels MM, Qualmann B. The actin nucleator Cobl is controlled by calcium and calmodulin. PLoS Biol (2015) 13: e1002233.

Izadi M, Schlobinski D, Lahr M, Schwintzer L, Qualmann B, Kessels MM. Cobl-like promotes actin filament formation and dendritic branching using only a single WH2 domain. J Cell Biol (2018) 217:211–230.

Izadi M, Seemann E, Schlobinski D, Schwintzer L, Qualmann B, Kessels MM. Functional interdependence of the actin nucleator Cobl and Cobl-like in dendritic arbor development. eLIFE (2021) 10: e67718.

Kessels MM, Qualmann B. Syndapin oligomers interconnect the machineries for endocytic vesicle formation and actin polymerization. J Biol Chem (2006) 281: 13285–13299.

Kessels MM, Engqvist-Goldstein AEY, Drubin DG, Qualmann B. Mammalian Abp1, a signal- responsive F-actin-binding protein, links the actin cytoskeleton to endocytosis via the GTPase dynamin. J Cell Biol (2001) 153: 351–366.

Koch D, Westermann M, Kessels MM, Qualmann B. Ultrastructural freeze-fracture immunolabeling identifies plasma membrane-localized syndapin II as a crucial factor in shaping caveolae. Histochem Cell Biol (2012) 138: 215–230.

Koch D, Spiwoks-Becker I, Sabanov V, Sinning A, Dugladze T, Stellmacher A, Ahuja R, Grimm J, Schüler S, Müller A, Angenstein F, Ahmed T, Diesler A, Moser M, Tom Dieck S, Spessert R, Boeckers TM, Fässler R, Hübner CA, Balschun D, Gloveli T, Kessels MM, Qualmann B. Proper synaptic vesicle formation and neuronal network activity critically rely on syndapin I. EMBO J (2011) 30: 4955–4969.

Liu M, Chen P, Hu HY, Ou-Yang DJ, Khushbu RA, Tan HL, Huang P, Chang S. Kinase gene fusions: roles and therapeutic value in progressive and refractory papillary thyroid cancer. J Cancer Res Clin Oncol (2021) 147: 323–337.

Marconi C, Canobbio I, Bozzi V, Pippucci T, Simonetti G, Melazzini F, Angori S, Martinelli G, Saglio G, Torti M, Pastan I, Seri M, Pecci A. 5’UTR point substitutions and N-terminal truncating mutations of ANKRD26 in acute myeloid leukemia. J Hematol Oncol (2017) 10: 18.

Morandi A, Plaza-Menacho I, Isacke CM (2011) RET in breast cancer: functional and therapeutic implications. Trends Mol Med 17: 149–157.

Mosavi LK, Cammett TJ, Desrosiers DC, Peng ZY. The ankyrin repeat as molecular architecture for protein recognition. Protein Sci (2004) 13: 1435–1448.

Mulligan LM. RET revisited: expanding the oncogenic portfolio. Nat Rev Cancer (2014) 14: 173–186

Nigg EA, Holland AJ. Once and only once: mechanisms of centriole duplication and their deregulation in disease. Nat Rev Mol Cell Biol (2018) 19: 297–312.

Noris P, Perrotta S, Seri M, Pecci A, Gnan C, Loffredo G, Pujol-Moix N, Zecca M, Scognamiglio F, De Rocco D, Punzo F, Melazzini F, Scianguetta S, Casale M, Marconi C, Pippucci T, Amendola G, Notarangelo LD, Klersy C, Civaschi E, Balduini CL, Savoia A. Mutations in ANKRD26 are responsible for a frequent form of inherited thrombocytopenia: analysis of 78 patients from 21 families. Blood (2011) 117: 6673–6680.

Pinyol, R Haeckel A, Ritter A, Qualmann B, Kessels MM. Regulation of N-WASP and the Arp2/3 complex by Abp1 controls neuronal morphology. PLoS One (2007) 2: e400.

Pippucci T, Savoia A, Perrotta S, Pujol-Moix N, Noris P, Castegnaro G, Pecci A, Gnan C, Punzo F, Marconi C, Gherardi S, Loffredo G, De Rocco D, Scianguetta S, Barozzi S, Magini P, Bozzi V, Dezzani L, Di Stazio M, Ferraro M, Perini G, Seri M, Balduini CL. Mutations in the 5’ UTR of ANKRD26, the ankirin repeat domain 26 gene, cause an autosomal-dominant form of inherited thrombocytopenia, THC2. Am J Hum Genet (2011) 88: 115–120.

Redfield SM, Mao J, Zhu H, He Z, Zhang X, Bigler SA, Zhou X. The C-terminal common to group 3 POTES (CtG3P): a newly discovered nucleolar marker associated with malignant progression and metastasis. Am J Cancer Res (2013) 3: 278–289.

Richardson DS, Lai AZ, Mulligan LM. RET ligand-induced internalization and its consequences for downstream signaling. Oncogene (2006) 25: 3206–3211.

Richardson DS, Gujral TS, Peng S, Asa SL, Mulligan LM. Transcript level modulates the inherent oncogenicity of RET/PTC oncoproteins. Cancer Res (2009) 69: 4861–4869.

Salvatore D, Santoro M, Schlumberger M. The importance of the RET gene in thyroid cancer and therapeutic implications. Nat Rev Endocrinol (2021) 17: 296–306.

Schneider K, Seemann E, Liebmann L, Ahuja R, Koch D, Westermann M, Hübner CA, Kessels MM, Qualmann B. ProSAP1 and membrane nanodomain-associated syndapin I promote postsynapse formation and function. J Cell Biol (2014) 205: 197–215.

Seemann E, Sun M, Krueger S, Tröger J, Hou W, Haag N, Schüler S, Westermann M, Huebner CA, Romeike B, Kessels MM, Qualmann B. Deciphering caveolar functions by syndapin III KO-mediated impairment of caveolar invagination. eLife (2017) 6: pii: e29854.

Shaw AT, Hsu PP, Awad MM, Engelman JA. Tyrosine kinase gene rearrangements in epithelial malignancies. Nat Rev Cancer (2013) 13: 772–787.

Staubitz JI, Musholt TJ, Schad A, Springer E, Lang H, Rajalingam K, Roth W, Hartmann N. ANKRD26-RET - A novel gene fusion involving RET in papillary thyroid carcinoma. Cancer Genet (2019) 238: 10–17.

Takahashi M, Cooper GM. Ret transforming gene encodes a fusion protein homologous to tyrosine kinases. Mol Cell Biol (1987) 7: 1378–1385.

Tanos BE, Yang HJ, Soni R, Wang WJ, Macaluso FP, Asara JM, Tsou MF. Centriole distal appendages promote membrane docking, leading to cilia initiation. Genes Dev (2013) 27: 163–168.

Tansey MG, Baloh RH, Milbrandt J, Johnson EM Jr. GFRalpha-mediated localization of RET to lipid rafts is required for effective downstream signaling, differentiation, and neuronal survival. Neuron (2000) 25: 611–623.

Wahlster L, Verboon JM, Ludwig LS, Black SC, Luo W, Garg K, Voit RA, Collins RL, Garimella K, Costello M, Chao KR, Goodrich JK, DiTroia SP, O’Donnell-Luria A, Talkowski ME, Michelson AD, Cantor AB, Sankaran VG. Familial thrombocytopenia due to a complex structural variant resulting in a WAC-ANKRD26 fusion transcript. J Exp Med (2021) 218: e20210444.

Wolf D, Hofbrucker-MacKenzie SA, Izadi M, Seemann E, Steiniger F, Schwintzer L, Koch D, Kessels MM, Qualmann B. Ankyrin repeat-containing N-Ank proteins shape cellular membranes. Nat Cell Biol (2019) 21: 1191–1205.

Yan H, Chen C, Chen H, Hong H, Huang Y, Ling K, Hu J, Wei Q. TALPID3 and ANKRD26 selectively orchestrate FBF1 localization and cilia gating. Nat Commun (2020) 11: 2196.

Zobel T, Brinkmann K, Koch N, Schneider K, Seemann E, Fleige A, Qualmann B, Kessels MM, Bogdan S. Cooperative functions of the two F-BAR proteins Cip4 and Nostrin in the regulation of E- cadherin in epithelial morphogenesis. J Cell Sci (2015) 128: 499–515.

